# Molecular crowding: impacts on the activity of the 10-23 DNAzyme

**DOI:** 10.64898/2026.06.30.735450

**Authors:** Nina Kirchgässler, Hannah Rosenbach, Ralf Biehl, Gerhard Steger, Richard Börner, Ingrid Span

## Abstract

The growing number of approved nucleic acid therapeutics illustrates the potential to treat diseases by targeting their genetic blueprints *in vivo*. The 10-23 DNAzyme is capable of cleaving a wide range of target RNA with high selectivity. However, its poor performance *in vivo* restricts its therapeutic application as gene silencing agent. Studies on ribozymes have shown that the crowded environment in cells and associated effects can impact ribozyme folding and thermostability, resulting in a change in activity. This opens up the question whether DNAzymes are also affected by molecular crowding. Here, we investigate the functional and structural influence of molecular crowding conditions on the 10-23 DNAzyme. The stability and activity of a PrP-specific 10-23 DNAzyme were examined in presence of PEG, dextran, and osmolytes. Our results indicate that osmolytes decrease DNAzyme activity in a concentration-dependent manner, while certain PEG and dextran concentrations promote activity. To rationalize our observations, we studied the cosolutes’ effect on physicochemical solution properties and the structure of the DNAzyme:RNA complex using FCS and SAXS. The data reveal that enhanced activity is observed under conditions where a combination of physiochemical properties matches an ‘optimum’ that seems to be dependent on the metal ion cofactor. Structural influence under such conditions is indicated less. We propose that a certain degree of molecular crowding is required to favor a state, which allows for higher catalytic turnover. In addition, we show that the requirement for magnesium and manganese as a cofactor remains unchanged under the conditions applied. Our work contributes to a better understanding of how the cellular environment affects DNAzyme structure and function.

## Introduction

In 1994, ***Breaker and Joyce (1994***) have found catalytically active, single-stranded DNA molecules by *in vitro* selection, called desoxyribozymes or DNAzymes. Adopting a three-dimensional structure allows DNAzymes to catalyze a broad spectrum of reactions (***Chandrasekar and Silverman, 2013; Hollenstein, 2015; Silverman, 2008; Zhang, 2018; Zhou et al., 2017***). Properties such as small in size, affordable in synthesis, diversity in modifications, and programmability made DNAzymes attract attention. Despite their diversity, RNA-cleaving DNAzymes in particular emerged to function as an alternative approach in nucleic acid therapeutics (***Karnati et al., 2014; Larcher et al., 2023; Thomas et al., 2021***).

Nowadays, one of the most prominent RNA-cleaving DNAzymes is the 10-23 DNAzyme (Dz, for reviews see ***Larcher et al., 2023; Rosenbach et al., 2020b; Yan et al., 2023***). Structurally, it consists of a 15 nt catalytic core flanked by two substrate binding arms, which may vary in length and sequence (usually 9–12 nt), but have to hybridize to the complementary RNA substrate of interest. To perform RNA cleavage, the Dz requires divalent cations (M^2+^), such as Mg^2+^ or Mn^2+^, and a purine-pyrimidine junction (5’–RY–3’) as cleavage site with an efficiency order of AU = GU ≥ GC ≫ AC (***Figure 1***A, B). Mechanistically, the Dz is predicted to perform cleavage of a phosphodiester bond via an acid-base mechanism (***Borggräfe et al., 2023; Santoro and Joyce, 1998***), similar to the 8-17 DNAzyme (***Cepeda-Plaza et al., 2018; Cepeda-Plaza and Peracchi, 2020; Cort’es-Guajardo et al., 2021***).

**Figure 1.**
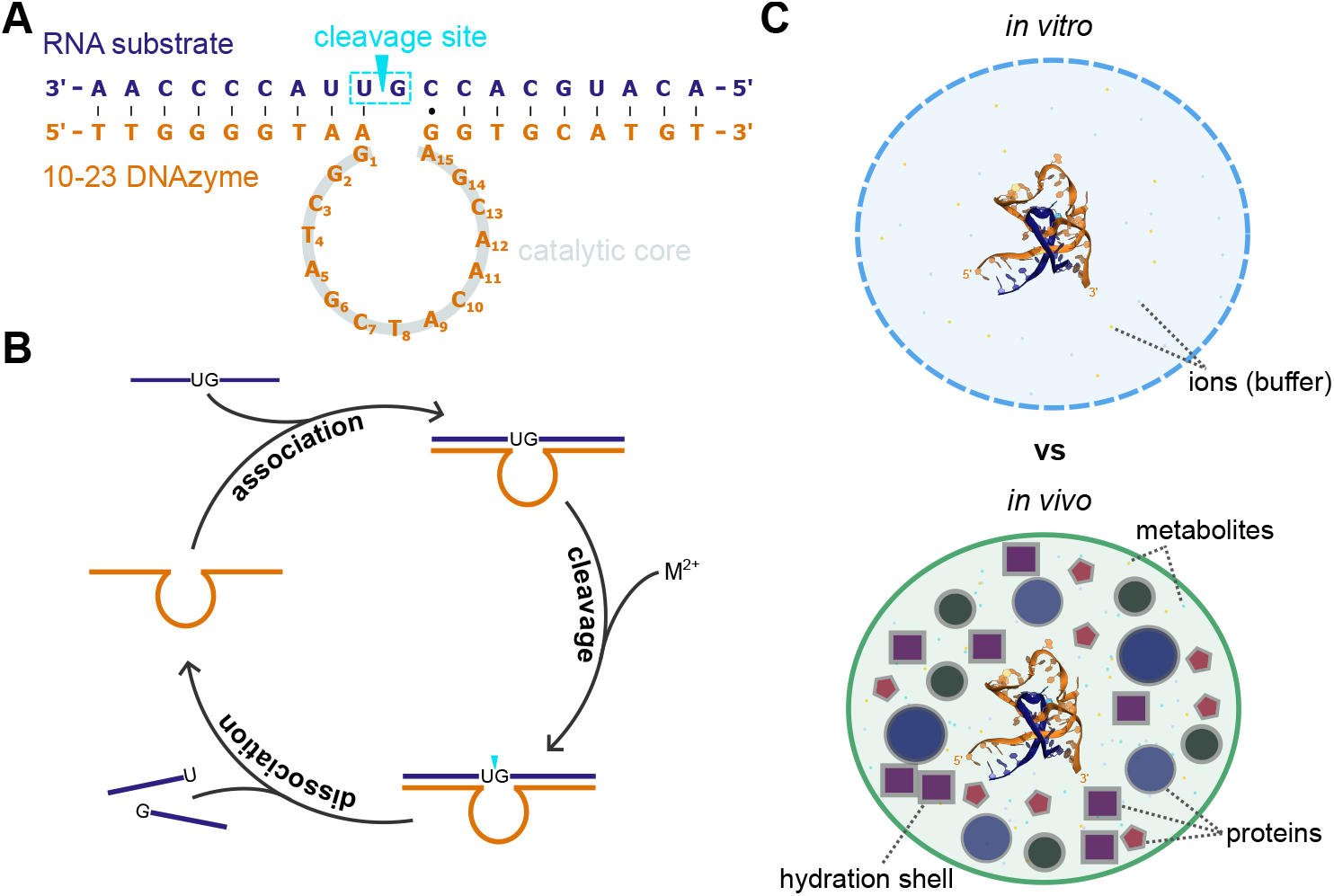
**A** Secondary structure of the 10-23 DNAzyme used in this study (Dz^WT^) in complex with its RNA substrate. **B** Catalytic cycle of the 10-23 DNAzyme, which illustrates cleavage under multiple turnover. After association of the Dz and RNA substrate by complementary base pairing, site-specific cleavage is induced depending on divalent cations M^2+^. Finally, the DNAzyme dissociates from the cleaved RNA to enter a new cycle. Modified after ***Rosenbach et al. (2020a***). **C** Simplified illustration of differences between analyses on the DNAzyme:RNA complex (PDB 7pdu, ***Borggräfe et al., 2021***) under *in vitro* and *in vivo* conditions.

Since their discovery, the spectrum of DNAzymes in terms of function and catalysis (e. g. phosphorylation, amide hydrolysis, Diels-Alder reaction) broadened immensely for *in vitro*-based applications (***Silverman, 2016; Zhang, 2018***). To this day, however, the use of DNAzymes in cells is limited due to a) poor understanding of the catalytic mechanism and b) low *in vivo* activity (***Etzkorn and Span, 2023; Garn and Renz, 2017***). Although first DNAzymes were reported that have entered clinical trials (e. g. ***Cho et al., 2013; Krug et al., 2017***), therapeutic success remains rare claiming several challenges: susceptibility to degradation by nucleases (***Xiao et al., 2023***), hampered accessibility of the target sequence by RNA folding (***Taylor et al., 2022***), low availability of free cofactors ([Mg^2+^] ≈ 0.5-1 mM, ***Victor et al. (2018***)), and activity modulating properties by monovalent ions (***Rosenbach et al., 2021***).

The latest modification studies on DNAzymes aim at addressing these limitations by partially modifying the backbone, sugar, and base of nucleotides, so-called xeno nucleic acids (XNAs). First attempts show progress in improved *in vivo* activities, thus providing a promising toolbox for the future (***Gerber et al., 2022; Taylor et al., 2022; Wang et al., 2021***). A certainly lasting challenge is differentiating RNA knockdown effects by targeted DNAzyme activity from RNase H-induced antisense effects (***Rivory et al., 2006; Taylor and Holliger, 2022; Zhou et al., 2022***), leaving room for further optimization and reliable controls.

For several years now, questions pointed towards potential effects on DNAzyme activity by molecular crowding in cells. In general, the term molecular crowding often summarizes effects resulting from macromolecular crowding, thus volume exclusion, confinement, and adsorption with thermodynamic and kinetic consequences. Since cells comprise a diversity of macromolecules, which generate a densely packed and heterogeneous environment, *in vivo* conditions differ clearly from experimentally controlled *in vitro* conditions (***Feig et al., 2017; Miyoshi and Sugimoto, 2008; Zhou, 2008; Zimmerman and Minton, 1993***, ***Figure 1***C). With this, several studies on double-stranded DNA and RNA, proteins, and ribozymes reported impacts on e. g. mobility, water reactivity, folding, and intermolecular interactions (***DasGupta, 2020; Fiorini et al., 2015; Kuznetsova et al., 2014; Nakano et al., 2014, 2008; Paudel et al., 2018***). Since the extent to which a biomolecule is affected by molecular crowding depends, amongst others, on its size and shape, this raises the question if DNAzymes are also affected by it.

In this study, we aim at simulating molecular crowding conditions *in vitro* using polyethylene glycol (PEG), dextran, and osmolytes as cosolutes and investigate their influence on DNAzyme function and structure. We hope that our results will provide first insights whether effects associated with molecular crowding interfere with DNAzyme activity that will help to better understand *in vivo* restrictions.

## Results

### Impact of crowding agents on DNAzyme activity and stability

To investigate effects of molecular crowding on the 10-23 DNAzyme, we used a variant that cleaves the mRNA of the PrP protein (for sequence see ***Figure 1***A; ***Victor et al., 2018***) and simulated crowded conditions *in vitro* using polymer-based cosolutes (PEG and dextran; ***DasGupta, 2020; Mardoum et al., 2018***), the PEG monomer EG as well as the osmolytes betaine and ectoine.

#### Oligonucleotides provide sufficient stability

To ensure sufficient stability of the oligonucleotides in presence of crowding, we performed assays in presence of the maximum cosolute concentration selected in this study, 0.18 g/mL, with subsequent analysis by urea-PAGE (***Figure 2***, ***Figure 2***—***figure Supplement 1***). All samples that contain the RNA substrate only (T) do not show additional bands besides the uncleaved molecule, which indicates construct stability under the conditions used for this analysis. In samples containing the RNA substrate and DNAzyme (TDz), bands indicate site-specific cleavage without further degradation of the fluorescent cleavage product. Taken together, the results indicate that all tested cosolutes and conditions provide oligonucleotide stability and were considered for further analysis.

**Figure 2.**
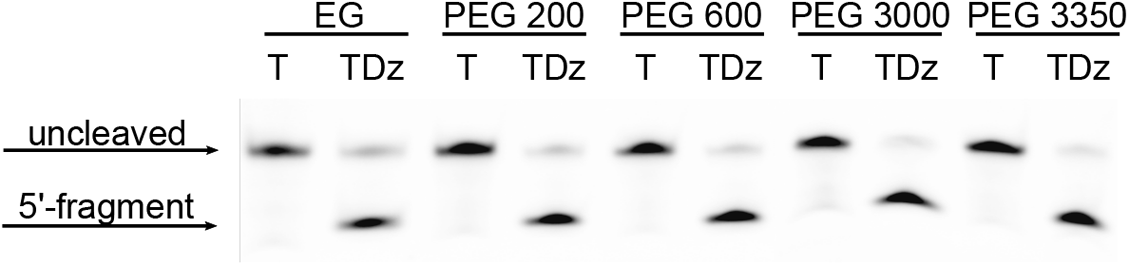
Urea-PAGE of samples after stability analysis in presence of 0.18 g/mL of cosolute. Conditions were analyzed in pairs of equally treated samples containing the RNA substrate only (T) and the DNAzyme:RNA complex (TDz). Assays were performed using the T^FAM^ and Dz^WT^ constructs (***Table 1***). Fluorescence detection and visualization of 6-FAM labeled RNA substrates and the 5’ cleavage fragment. **Figure 2—figure supplement 1**. Urea-PAGE of samples after stability analysis in presence of 0.18 g/ml of the remaining cosolutes

#### Crowded conditions alter DNAzyme activity

As a next step, we investigated the effect of a large variety of crowded conditions on DNAzyme activity and performed FRET-based cleavage assays under single-turnover conditions with cosolute concentrations of 0.03, 0.07, 0.11, 0.15 and 0.18 g/mL. Since Dz-mediated cleavage can be performed in presence of different metal ions, that demonstrated diverging impact at physiological relevant conditions in previous studies (0.5 mM M^2+^, 100 mM NaCl; ***Rosenbach et al., 2021***), assays were performed in presence of Mg^2+^ (***Figure 3***, ***Figure 3***—***figure Supplement 1***) and Mn^2+^ (***Figure 3***—***figure Supplement 2***).

**Table 1.**
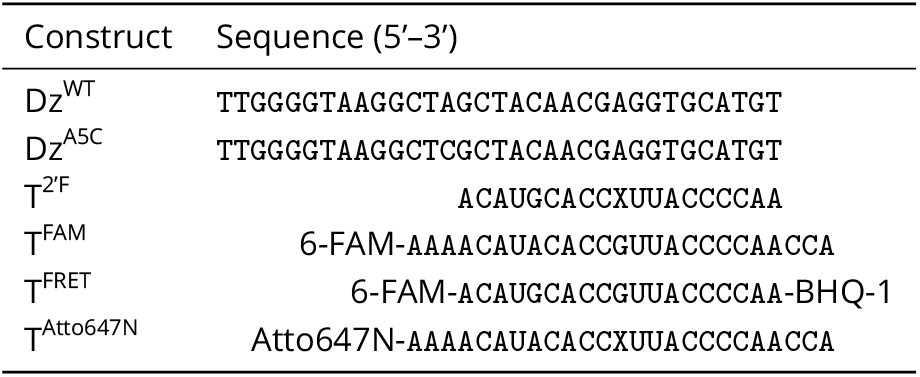
Sequences of Dz and RNA substrates used in this study. X=2’-F-dG.

**Figure 3.**
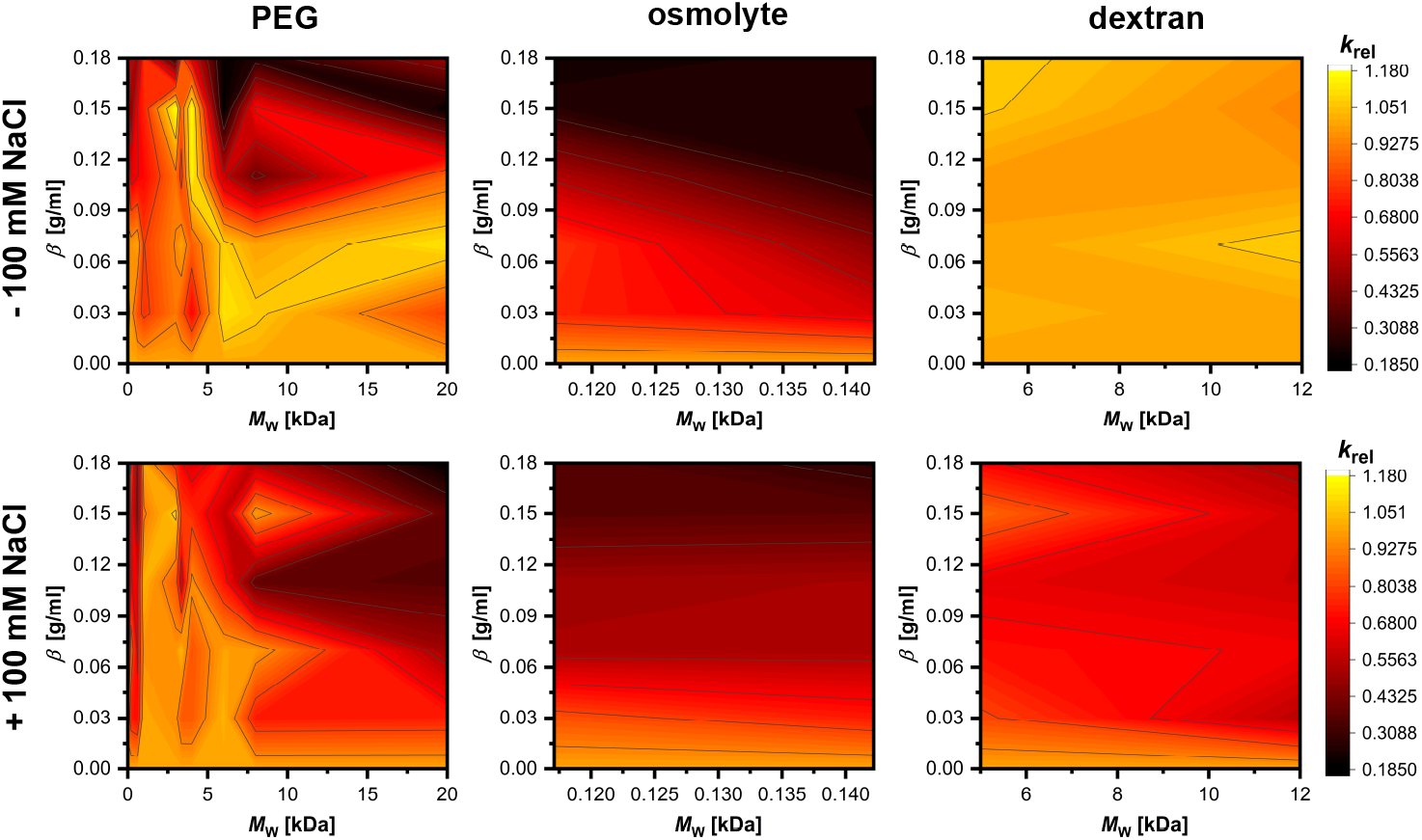
Effects of crowded conditions on Dz activity in presence and absence of 100 mM NaCl at 0.5 mM MgCl_2_. Dz activity was determined in FRET-based activity assays in presence of PEG, osmolyte or dextran. Relative rate constants *k*_rel,*i*_ ((2)) are based on determined rate constants *k*_obs,*n*=14_ = (0.0048 ± 0.0008) s^−1^ and *k*_obs,*n*=14_ = (0.0095 ± 0.0017) s^−1^ with and without 100 mM NaCl, respectively. **Figure 3—figure supplement 1**. Selection of cosolute concentrations that included conditions with promoting properties on Dz activity at 0.5 mM MgCl_2_ **Figure 3—figure supplement 2**. Effects of crowded conditions on Dz activity in presence and absence of 100 mM NaCl at 0.5 mM MnCl_2_ **Figure 3—figure supplement 3**. Dz activity depending on MgCl_2_ concentration

From the activity assays, we are able to observe different impacts on Dz activtiy depending on cosolute type, size, and concentration. In most cases, our data show that increasing cosolute concentrations lead to a decrease in Dz activity. Interestingly, in presence of certain conditions containing PEG and dextran we observe promoting effects on activity (*k*_rel_ > 1.10). Considering Mg^2+^-induced cleavage, favoring conditions include PEG variants < 6 kDa and concentrations of 0.11 and 0.15 g/mL in absence of NaCl (***Figure 3***). In presence of NaCl, promoting conditions diverge slightly and include cosolute concentrations of 0.07 and 0.15 g/mL (***Figure 3***—***figure Supplement 1***). Here, however, the extent, to which Dz activity is increased, is less than in absence of NaCl. *k*_rel_ has its maximum value at 0.07 g/ml of PEG 3350 of 1.07. In presence of Mn^2+^, promoting conditions shift to cosolute concentrations ≤ 0.07 g/mL, but appear to be similar for approaches with and without NaCl (***Figure 3***—***figure Supplement 2A***,***B***). In general, PEG variants with > 6 kDa result in decreased Dz-activity at concentrations ≥ 0.11 g/mL. Crowding simulated by the osmolytes betaine and ectoine result in a concentration-dependent decrease in Dz activity regardless of the respective cofactor, but with greater impact by ectoine (***Figure 3***, ***Figure 3***—***figure Supplement 2A***). Samples that contain dextran, on the other hand, show maintained or even enhanced activity in absence of NaCl. Promoting effects on activity are only observed in presence of dex 12000 and for Mn^2+^-induced cleavage at concentrations of 0.07 and 0.15 g/mL (Figure 2). Addition of NaCl result in fluctuations in activity with an overall greater impact by dex 12000, and in particular for Mg^2+^-induced catalysis.

#### Cofactor requirement and competition with ATP

Previous studies on ribozymes presented a decrease in cofactor requirement in presence of cosolutes that facilitated catalysis at near physiological concentrations (***DasGupta et al., 2023; Paudel et al., 2018***). Since low cofactor availability and competition with ATP for free Mg^2+^ are suggested to challenge DNAzyme approaches *in vivo*, we questioned the influence of molecular crowding on the Dz’s cofactor requirement (***Figure 4***, ***Figure 3***—***figure Supplement 2D***) and ability to compete with ATP (***Figure 3***—***figure Supplement 1***). Assays were performed in presence of crowded conditions that promoted Dz activity the most. The cofactor requirement was analyzed using FRET-based activity assays using MgCl_2_ and MnCl_2_ concentrations of 0 − 5 mM and 0 − 0.6 mM, from which we observe that a crowded environment did not evidently change cofactor requirement, at least for the conditions tested here. In absence as well as presence of NaCl, crowded conditions indicate no clear change in requirement for Mg^2+^. Only presence of 0.15 g/mL of PEG 4000 demonstrate a slight increase in activity at MgCl_2_ concentrations ≤ 0.5 mM compared to the buffer only. With Mn^2+^ as cofactor, presence of cosolutes results in an increases in activity between 0.4 − 0.6 mM MnCl_2_, below, there is no difference compared to the buffer only sample.

**Figure 4.**
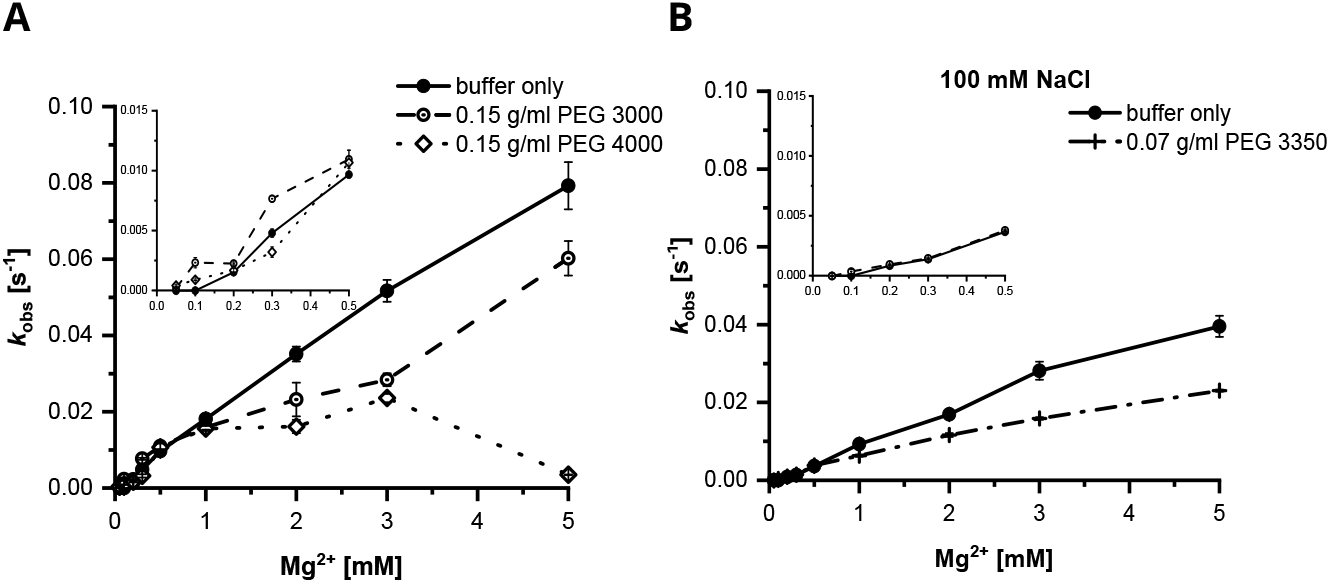
Cofactor-dependent rate constants in presence of crowded conditions with most activity enhancing properties: 0.15 g/ml of PEG 3000 and PEG 4000 in absence (**A**) and 0.07 g/ml of PEG 3350 in presence of 100 mM NaCl (**B**). Measurements were performed in a range of 0.05 − 5 mMMgCl_2_. **Figure 4—figure supplement 1**. Dz activity in presence of crowding and competition with ATP at 0.5 mM MgCl_2_ and 100 mM NaCl

ATP-competition assays revealed an ATP-dependent decrease in Dz activity in absence of crowding (***Figure 4***—***figure Supplement 1***), supporting the data published by ***Victor et al. (2018***). The same tendency also applies for all selected crowded conditions. Interestingly, Dz activity is less impacted by ATP when a crowded environment is mimicked by larger PEG. Here, activity at 0.18 g/mL PEG 20000 is hampered less compared to measurements with 0.15 g/mL PEG 600. Thus, we suggested that the extent of changes in solution properties determines diffusion-dependent binding of Mg^2+^ to either ATP or the Dz:RNA complex.

#### Activity in presence of osmolytes depending on ionic strength

Our data show that increasing concentrations of osmolytes enhance activity impairment. In absence of NaCl and presence of 0.18 g/mL betaine or ectoine, Dz activity is decreased to 22 − 26 % (0.5 mM MgCl_2_) or to 26 % (betaine) or rather 7 % (ectoine, 0.5 mM MnCl_2_) of the activity in absence of crowding. Assays in presence of 100 mM NaCl indicate, however, that the extent, to which osmolytes hamper Dz activity, changes (***Figure 3***, ***Figure 3***—***figure Supplement 2***). Thus, we questioned the effect’s dependence on ionic strength. FRET-based activity assays were performed in presence of NaCl concentrations ranging from 0 − 750 mM (***Figure 5***). To evaluate changes depending on ionic strength, relative rate constants ((2)) were determined in relation to Dz activity in absence of crowding and NaCl (*k*_obs, 0_).

**Figure 5.**
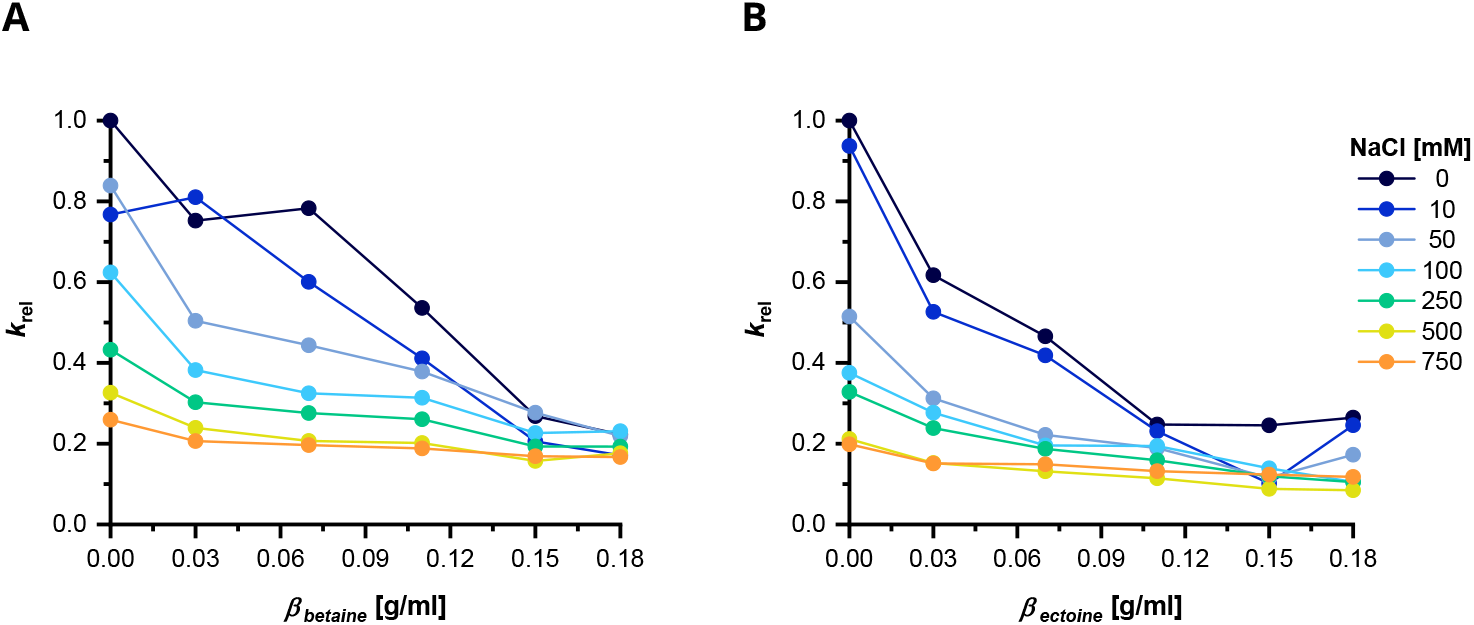
Effect of osmolytes on Dz activity depending on ionic strength. Analyses were performed by FRET-based cleavage assays at 0.5 mM MgCl_2_. Relative Dz activity in presence of different concentrations of betaine (**A**) or ectoine (**B**). *k*_rel_ was calculated based on *k*_obs, 0_ = 0.009 58 s^−1^ for experiments with ectoine and *k*_obs, 0_ = 0.007 35 s^−1^ with betaine.

Here, elevated levels of monovalent ions seem to lower the concentration-dependent effect of both osmolytes on catalysis. Sole presence of Na^+^-ions resulted in a decrease in activity of 80 % at 750 mM NaCl. Assays in presence of betaine (***Figure 5A***) indicated a gradual reduction in Dz activity with increasing NaCl concentration. Only 0.07 g/mL betaine in absence and 0.03 g/mL in presence of 10 mM NaCl seem to have less impact. Samples containing ectoine, however, not only resulted in a clear decrease in activity of −40 % at 0.03 g/mL of ectoine (0 mM NaCl, ***Figure 5B***), but also indicated an even greater impact, when NaCl is increased from 10 mM to 50 mM.

### Complex size

So far, we were able to observe that Dz activity is altered in presence of crowded conditions, but without a reduction or change in cofactor requirement. Since crowding studies in general stated altered solution properties as one central cause with thermodynamic and kinetic consequences, we further pursued investigations that target physiochemical solution properties and the Dz:RNA complex’s diffusion behaviour and size, of which the latter are intended to help rationalize our observations with structural data.

#### Physiochemical properties of crowding solutions

We determined the physiochemical solution properties in absence of the Dz:RNA complex and aimed at investigating changes within the Dz’s environment by crowding. Here, we performed density (***Figure 6A***), viscosity (***Figure 6B, Figure 6***—***table Supplement 1***), and pH measurements (***Figure 6C***). In the latter, conditions were selected to cover all cosolute types and different degrees of Dz activity. We observed a strong increase of the viscosity at high concentrations of large cosolutes, while the density and pH were less affected. Thus, we propose that viscosity-related effects potentially have a greater impact on Dz-mediated cleavage.

**Figure 6.**
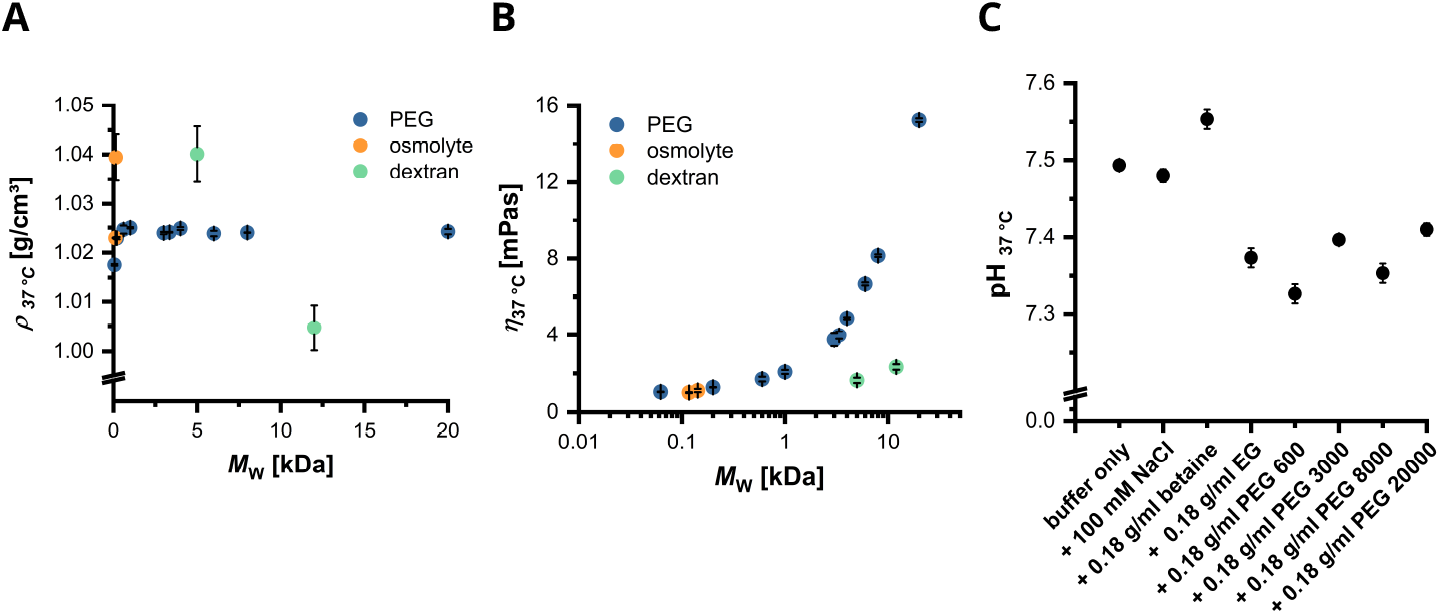
Changes in physiochemical solution properties in presence of cosolutes. **A** Density *ρ* of crowded solutions at 0.18 g/ml of cosolute and 37 °C. *ρ*_buffer only, 37 °C_ = (0.9952 ± 0.0003) g/cm^3^. **B** Dynamic viscosity *η* of crowded solutions at 0.18 g/ml of cosolute and 37 °C. *η*_buffer only, 37 °C_ = (0.7025 ± 0.0015) mPa·s. **C** pH of crowded solutions at 37 °C. Conditions were selected to cover different degrees of Dz activity. **Figure 6—figure supplement 1**. Hydrodynamic radius (*r*_h_, **A**), radius of gyration (*r*_g_, **B**), and theoretical, average distance between cosolutes (*D*, **C**) depending on *M*_W_ and concentration **Figure 6—table supplement 1**. Density *ρ* and dynamic viscosity *η* at 37 °C of additional samples used in FCS measurements with cosolute concentrations less than 0.18 g/ml **Figure 6—table supplement 2**. *D* of all used PEG variants at a concentration range of 0.03 to 0.18 g/mL

Density measurements showed a maximum shift of 5 % (Δ*ρ* = 0.045 g/cm^3^) at 0.18 g/mL of dextran 5000 compared to the buffer-only sample (*ρ*_buffer, 37 °C_ = (0.995 19 ± 0.000 32) g/cm^3^ (***Figure 6A***). In contrast, the solution’s viscosity changed profoundly with increasing size of cosolute (***Figure 6B***). Here, 0.18 g/mL of PEG 20000 increased the viscosity to the largest extent to (15.2610 ± 0.0929) mPa·s (21-fold) compared to the buffer only sample (*η*_buffer, 37 °C_ = (0.7025 ± 0.0015) mPa·s, ***Figure 6B***). Measurement of pH did not result in considerable changes with maximum shifts of ΔpH = −0.17 (0.18 g/mL PEG 600) and ΔpH = +0.05 (0.18 g/mL betaine, ***Figure 6C***).

#### Cosolute size and concentration limit accessible volume

As an increase in cosolute concentration and size not only affects the viscosity, but also determines the volume available between two cosolute molecules, we calculated the theoretical, average distance between them (cavity size, *D*) under the assumption of a hard sphere model. With this, we aim at investigating the extent, to which steric influence may impact Dz:RNA complex mobility and size. Determining *D* requires information on the hydrodynamic radius *r*_h_ (***Figure 6***—***figure Supplement 1A***, ***Equation 6, Equation 7***) and radius of gyration *r*_g_ (***Figure 6***—***figure Supplement 1B***, ***Equation 5***), which have been calculated based on previous studies. Since PEG is one of the best characterized and most frequently used polymers for crowding analyses, *D* was determined only for the variants used here (***Figure 6***—***figure Supplement 1C***, ***Equation 8, Equation 9***, and ***Equation 10***, ***Figure 6***—***table Supplement 2***). In this study, *D* was assumed to be 100 nm for the buffer only sample. ***Borggräfe et al. (2021***) published a first structure of the precatalytic Dz:RNA complex using the same Dz, except for one base substitution within the catalytic core, and presented a size of approximately 50 Å in diameter. Taking this into account, *D* does not reach the proposed complex size for the most tested cosolutes and respective concentrations. However, the size is approximated, which includes *D* between five and six nm, or even smaller at ≥ 0.11 g/mL of EG, 0.18 g/mL of PEG 200, PEG 600, PEG 1000, and PEG 6000, ≥ 0.15 g/mL of PEG 8000, and ≥ 0.11 g/mL of PEG 20000. In general, the calculated values of *D* cover a size range of ~ 4.4 − 11.2 nm.

Since structural information by NMR did not include a crowded environment, we performed fluorescence correlation spectroscopy (FCS) and small-angle X-ray scattering (SAXS) experiments to obtain and compare new structural insights in presence of cosolutes. Due to a lack of information to calculate *D* for betaine, ectoine, and dex 12000, we assumed that *D*(0.18 g/mL betaine) = *D*(0.18 g/mL ectoine) = *D*(0.18 g/mL EG) and *D*(0.18 g/mL dextrane 1200) = *D*(0.11 g/mL PEG 8000).

#### Changes in solution properties impact Dz:RNA complex size

Diffusion analyses by FCS were carried out to determine the constructs’ diffusion time *τ*_D_ (***Figure 7***— ***figure Supplement 1***, ***Equation 3***) and consequently the hydrodynamic radius *r*_h_ (***Figure 7, Equation 4***). The data indicate a dependence between *r*_h_ and *τ*_D_ for T and TDz with cosolute size (***Figure 7E***,***Figure 7***—***figure Supplement 1C***), concentration, and eventually associated solution properties *η* (***Figure 7D, Figure 7***—***figure Supplement 1B***) and *D* (***Figure 7C, Figure 7***—***figure Supplement 1A***).

**Figure 7.**
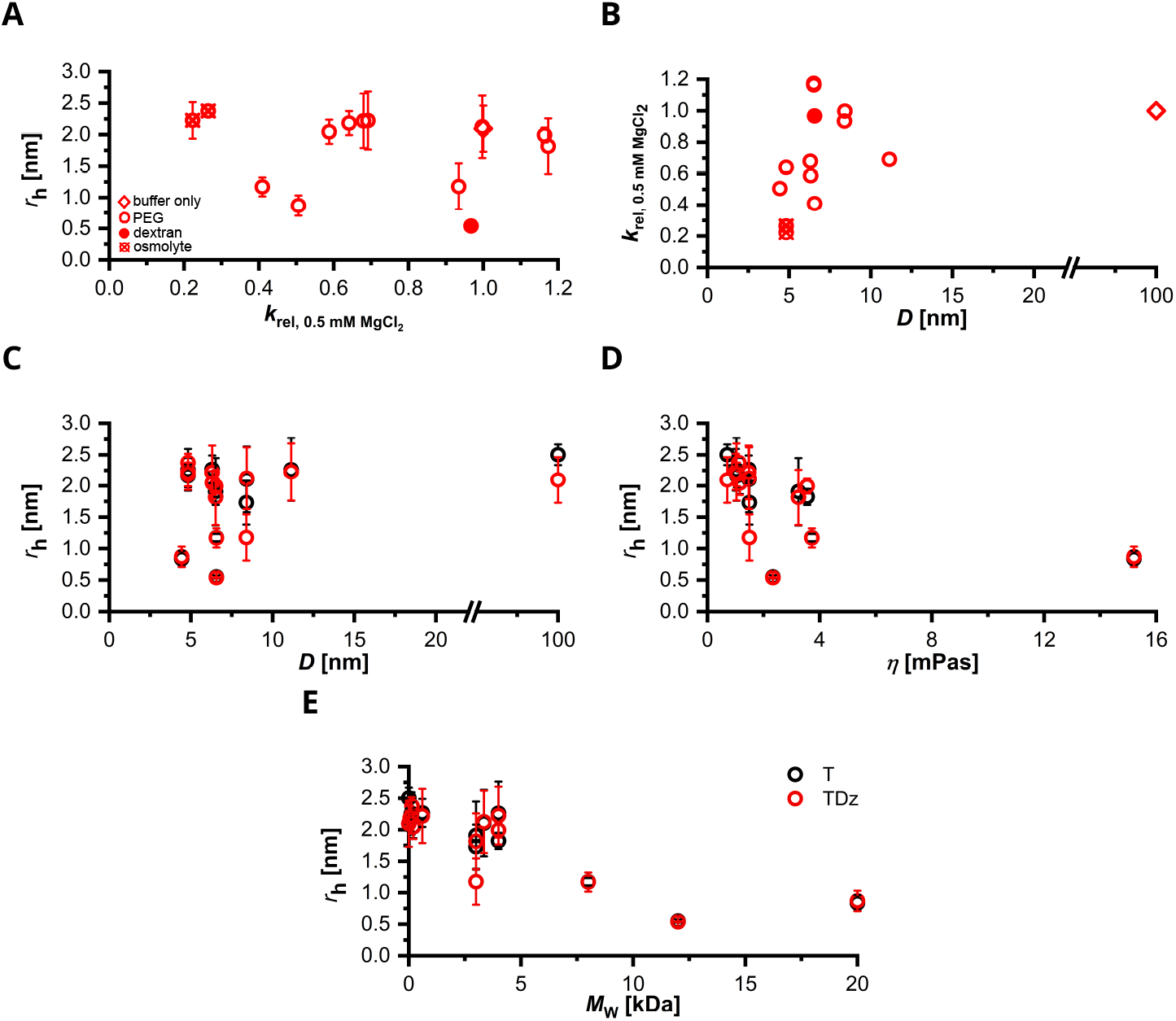
Hydrodynamic radius *r*_h_ of RNA substrate (T) and Dz:RNA complex (TDz) in absence and presence of cosolutes determined by FCS. **A** *r*_h_[nm] of TDz in comparison to Dz activity as *k*_rel_ ((2)) at 0.5 mM MgCl2_2_ (*k*_obs,*n*=14_ = (0.0095 ± 0.0017) s^−1^). **B** Dz activity as *k*_rel_ at 0.5 mM MgCl2_2_ in relation to cavity size *D*. **C** *r*_h_ depending on cavity size *D*. **D** *r*_h_ depending on solution viscosity at 37 °C. **E** *r*_h_ depending on *M*_W_ of the respective cosolute. **Figure 7—figure supplement 1**. Diffusion time *τ*_D_ of RNA substrate (T) and Dz:RNA complex (TDz) in absence and presence of cosolutes determined by FCS. **Figure 7—figure supplement 2**. Comparison of the Dz:RNA complex’s hydrodynamic radius *r*_h_, *D*, and relative Dz activity for Mn^2+^-induced RNA cleavage.

In absence of crowding, the Dz:RNA complex is slightly smaller than the RNA substrate alone (*r*_h, TDz_ = (2.09 ± 0.37) nm, *r*_h, T_ = (2.50 ± 0.17) nm). Here, *τ*_D_ of T is (0.193 ± 0.025) ms and of TDz is (0.161 ± 0.029) ms (***Figure 7***—***figure Supplement 1***). In correlation with cosolute size and concentration, *τ*_D_ increases for T and TDz to a similar extent and reaches a maximum at 0.18 g/mL PEG 20000 (*τ*_D, T_ = (1.380 ± 0.108) ms, *τ*_D, TDz_ = (1.429 ± 0.150) ms). Under crowded conditions that contain PEG as cosolute, our data do not indicate a direct link between changes in *r*_h_ and solution properties such as *D* (***Figure 7C***) and *η* (***Figure 7D***), which makes us assume that further environmental factors impact Dz:RNA complex size. On the contrary, presence of betaine and ectoine as well as EG slightly increases *r*_h, TDz_ to a maximum of *r*_h, TDz_ = (2.38 ± 0.09) nm at 0.18 g/mL of ectoine. Interestingly, of all tested samples, 0.18 g/mL of dex 12000 (*η* = 2.33 mPa·s) led to the strongest compaction of T and TDz (*r*_h, T_ = (0.55 ± 0.03) nm, *r*_h, TDz_ = (0.54 ± 0.01) nm), although Dz activity was not affected significantly (*k*_rel_ = 0.97, ***Figure 7A***). Even though FCS have been measured in absence of Mg^2+^ or Mn^2+^, conditions that demonstrated promoting effects on Mg^2+^-induced Dz activity (*k*_rel_ > 1.10, ***Figure 3***) resulted in *r*_h, TDz_ between 1.81 and 2.0 nm (***Figure 7A, Figure 7B***). Here, *D* is 6.51 nm. In presence of NaCl, *D* of favoring conditions shifts slightly and ranges between 6.51 and 8.41 nm. In contrast, conditions that promoted Dz activity with Mn^2+^ as cofactor (*k*_rel_ ≥ 1.10, ***Figure 3***—***figure Supplement 2***) demonstrate a *r*_h, TDz_ between 2.12 and 2.22 nm and include *D* of 8.40 and 11.1 nm (***Figure 7***—***figure Supplement 2***). In presence of NaCl, 0.03 g/ml of PEG 4000 as the most promoting condition (*k*_rel_ = 1.13) underlines an apparent preference of *D* of 11.1 nm with Mn^2+^ as cofactor. Overall, the data indicate that a hydrodynamic radius of ~ 1 − 2.22 nm applies for different degrees of Dz activity (*k*_rel_ = 0.4-1.17). In this regard, it is interesting that a *r*_h, TDz_ within a range of ~ 1.81 − 2.22 nm can correspond to a *k*_rel_ < 0.3 as well as > 1.0 for Mg^2+^ and Mn^2+^. Additionally, we observe that crowded conditions that cause the Dz:RNA complex to compact and result in *r*_h, TDz_ of < 1 nm restrict activity, which mainly occurs in presence of cosolutes with *M*_W_ > 5 kDa.

Aiming for additional information on Dz:RNA complex size and shape, we performed SAXS measurements (***Figure 8***). Analyses in presence of MgCl_2_ were further carried out using the A5C variant (Dz^A5C^, Table ***Table 1***). Substitution of the catalytic core residue A5 disrupts the palindromic sequence within the catalytic core and prevents homodimer formation (***Borggräfe et al., 2021***). A comparison of both sequences will indicate whether samples contain complex species differing from the active complex that influence model predictions. We observe an elongated shape for the Dz:RNA complex with only small deviations between samples in absence and presence of crowding. In addition, our data reveal a slightly smaller size for the Dz:RNA complex containing the A5C variant than the wild type DNAzyme.

**Figure 8.**
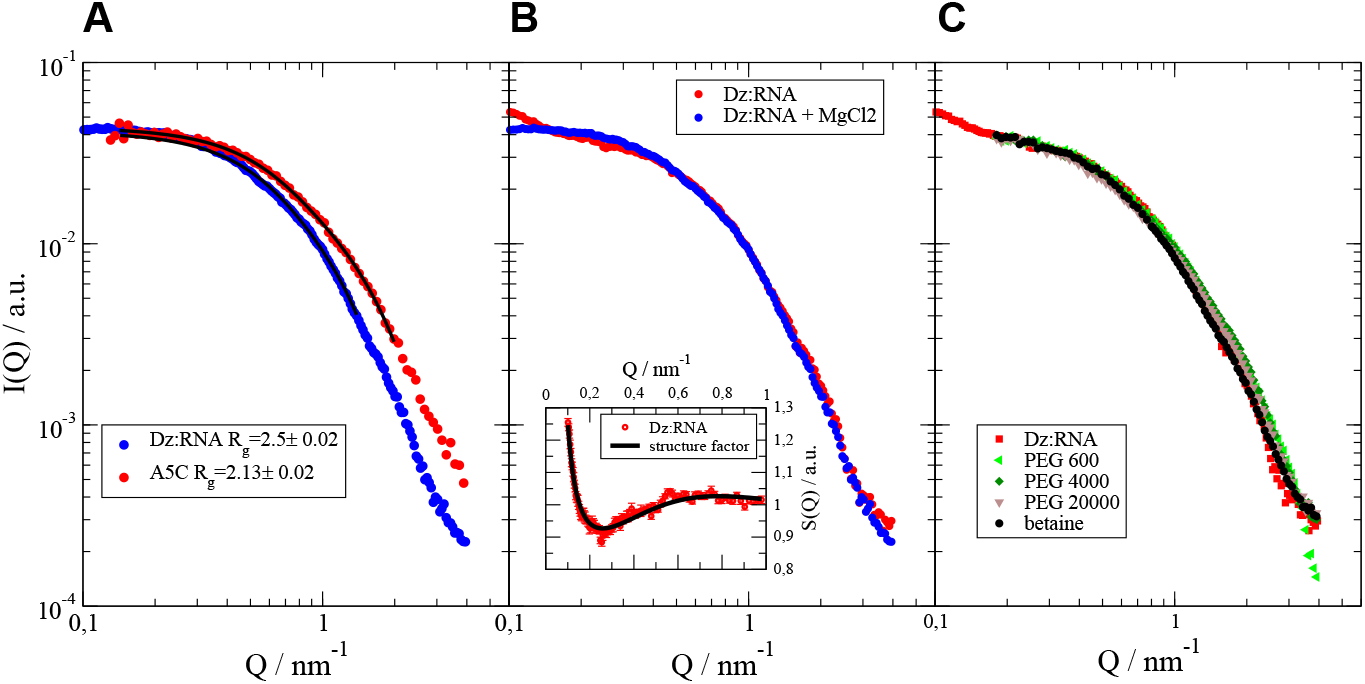
SAXS scattering intensities *I*(*Q*) in dependence upon scattering vector *Q*. **A** Comparison of the Dz:RNA complex that either contains the Dz^WT^ (blue) or the Dz^A5C^ variant (red) in presence of 5 mM MgCl_2_. Black lines show a fit with an ellipsoid of revolution form factor. **B** Comparison of the Dz^WT^:RNA complex in absence (blue) and presence (red) of 5 mM MgCl_2_. The inset shows the structure factor responsible for the deviation with and without MgCl_2_. **C** Comparison of Dz^WT^:RNA complex in absence (red) and presence of 0.18 g/ml of the respective cosolute. PEG is known to be nearly matched in water that it is not seen in SAXS (***Thiyagarajan et al., 1995***).

In presence of 5 mM MgCl_2_, the Dz^A5C^:RNA complex shows a more compact state (*r*_*g*_ = 2.1 nm) than the Dz^WT^-containing construct (*r*_*g*_ = 2.5 nm, ***Figure 8A***). A ratio of *r*_*g*_/*r*_*h*_ ≈ 1.2 compared to *r*_*g*_/*r*_*h*_ ≈ 0.77 for a solid sphere indicates already an elongated shape. Assuming an ellipsoidal shape, the Perrin friction factor indicates an aspect ratio ~ 5 (***Perrin, 1934***). Fitting an ellipsoidal formfactor to the scattering data in ***Figure 8A*** result in half axes *r*_*a*_ = (5.00 ± 0.05) nm, *r*_*b*_ = (1.08 ± 0.01) nm for the Dz^A5C^:RNA complex and half axes *r*_*a*_ = (5.37 ± 0.05) nm, *r*_*b*_ = (1.49 ± 0.01) nm for the Dz^WT^:RNA complex, which supports the assumption of an elongated shape. While a diameter of 2*r*_*b*_ equals roughly the diameter of DNA with 1.8 − 2.3 nm, the total length of 2*r*_*a*_ can only be reached with a strongly stretched configuration of 19 bp. Nevertheless, an elongated structure of the complex can be assumed.

In ***Figure 8B***, the Dz^WT^:RNA complex in absence of MgCl_2_ presents a deviation from samples with MgCl_2_, which we observe from a higher scattering intensity *I*(*Q*) at lowest scattering vector *Q* and lower values near 0.2 nm^−1^. This deviation is characteristic for a structure factor due to interparticle interactions with a short-range attraction and long-range repulsion. The structure factor *S*(*Q*) for the Dz^WT^:RNA complex can be determined using the complex’s sample with 5 mM MgCl_2_ as a formfactor (see ***Figure 8A*** for comparison) and dividing the measured *I*(*Q*) by this formfactor *F* (*Q*) according to *I*(*Q*) = *S*(*Q*)*F* (*Q*). The extracted structure factor *S*(*Q*) (see inset in ***Figure 8B***) shows good agreement with a two Yukawa structure factor with a short range attractive and long-range repulsive component. Exchange of Na^+^ at the complex by Mg^2+^ seems to reduce attraction and repulsion. ***Figure 8C*** compares the scattering patterns of the Dz^WT^:RNA complex with different cosolutes in the solution. In presence of betaine, the complex shows only a small deviation to the sample without crowding. In presence of PEG, smaller deviations for *Q* > 1 nm^−1^ might indicate that conformational changes are much smaller than the difference between the Dz^WT^:RNA and Dz^A5C^:RNA complex in ***Figure 8A***.

## Discussion

High specificity, low cost in synthesis, and flexibility in design made DNAzymes as molecular tools gain attention, of which especially RNA-cleaving variants were considered promising to function as gene silencing agents. However, low *in vivo* activity remains a major challenge in cell-based Dz applications, in which effects by molecular crowding have been proposed to contribute (***DasGupta, 2020; Paudel et al., 2018; Paudel and Rueda, 2014; Rosenbach et al., 2021; Victor et al., 2018***). Studies on ribozymes revealed changes in activity, cofactor requirement, and folding, in which crowding favored their activity at physiological relevant Mg^2+^ concentrations (***Nakano et al., 2008***) and promoted formation and stabilization of a compact and active conformation (***Paudel et al., 2018***). Thus, shedding light on possible effects by molecular crowding on the 10-23 DNAzyme is of great relevance for a fundamental understanding of Dz catalysis and for improving Dz by introducing chemical modifications. In this study we focus on functional analyses, changes in physiochemical solution properties, and the structure and shape of the Dz:RNA complex. Crowded conditions were mimicked using different concentrations of polymers PEG and dextran, EG, and osmolytes betaine and ectoine. With this approach we aim at covering different molecule properties and degrees of molecular crowding to achieve conditions that resemble a cell’s divers composition.

In comparison to polymers PEG and dextran, our activity assays demonstrate a concentration-dependent impact on Dz activity in presence of betaine and ectoine. The two exhibit zwitterionic properties with strong water binding at neutral pH (***Bownik and Stę pniewska, 2016; Kurz, 2008***). As the presence of both did not significantly change the density (***Figure 6A***), viscosity (***Figure 6B***), and pH (***Figure 6C***), we propose altered water activity as possible cause for activity impairment. Since both molecules are small in size, they are assumed less likely to influence cleavage by volume exclusion. Former *in vitro* studies using betaine revealed destabilization of nucleic acids depending on GC-content, with preferred accumulation around single-stranded DNA (***Hong et al., 2004; Swinefus et al., 2013; Vasudevamurthy et al., 2009***). Since the Dz’s catalytic core provides unpaired GC nucleotides exposed to the surface, betaine may be able to interact at multiple sites. Similar to its derivative hydroxyectoine, ectoine is suggested to prefer negatively charged surfaces leading, if bound, to strong, unspecific interactions, which occur within the first hydration shell (***Meyer et al., 2017; Oprzeska-Zingrebe et al., 2018***). Presence of both molecules resulted in a slight increase in *r*_h_ for the Dz:RNA complex (+0.132 nm at 0.18 g/mL betaine, +0.282 nm at 0.18 g/mL ectoine; ***Figure 7***). This in combination with their interaction potential makes consider complex dehydration and consequently elongation as possible cause. Due to their zwitterionic character and concentration-dependent impact, we questioned the effect’s dependence on ionic strength (***Figure 5***). From activity assays in presence of different Na^+^ concentrations we observe less impact by the two osmolytes at elevated Na^+^ levels. As Na^+^ ions are proposed to compete with Mg^2+^ for binding sites (***Rosenbach et al., 2021***), their sole presence is sufficient to diminish Dz activity.

In contrast, our activity assays in presence of PEG and dextran indicate that single crowded conditions promote activity (***Figure 3***), which matches observations in ribozyme-based analyses (***DasGupta et al., 2023; Paudel et al., 2018; Paudel and Rueda, 2014***). Presence of the monomer EG, on the other hand, led to a concentration-dependent decrease in Dz activity. Previous studies on nucleic acids and ribozymes also assigned osmolytic properties and effects to EG, PEG 200, and PEG 600, by altering water activity (***Adams and Znosko, 2019; DasGupta, 2020; DasGupta et al., 2023; Ghosh et al., 2020***). Based on our activity measurements, we assume similar effects for Mg^2+^-(***Figure 3***) and Mn^2+^-induced (***Figure 3***—***figure Supplement 1***) cleavage. Crowded conditions that promote Dz activity contain PEG variants with *M*_W_ > 1 kDa, of which most solutions include PEG 3000 and PEG 4000. However, preferred conditions seem to differ between Mg^2+^- and Mn^2+^-induced catalysis. Assays in presence of Mg^2+^ highlight an enhanced activity at 0.11 − 0.15 g/mL of cosolute (***Figure 3***), whereas the range shifts to 0.03 − 0.07 g/mL in presence of Mn^2+^ (***Figure 3***— ***figure Supplement 1***). But in presence of NaCl, lower cosolute concentrations seem to be beneficial, as 0.07 g/ml of PEG 3350 provided the most promoting surrounding with Mg^2+^ as cofactor and 0.03 g/ml of PEG 4000 with Mn^2+^. Depending on the respective cofactor, ***Rosenbach et al. (2021***) already predicted an altered interplay between ions, which the authors assigned to the metal ion’s properties. Our activity data also imply that effects by molecular crowding change depending on the cofactor. Analyses, for example, by ***Nakano et al. (2008***) demonstrated a lowered cofactor requirement for the hammerhead ribozyme in presence of crowding. Our activity assays, however, show that even the most promoting crowded conditions did not change the Dz’s cofactor requirement (***Figure 3***—***figure Supplement 2***, ***Figure 3***—***figure Supplement 1***). Although ATP competition assays indicate that altered solution properties in presence of larger PEG molecules improve Mg^2+^-binding to Dz, limited cofactor availability remains a considerable *in vivo* challenge (***Figure 4***—***figure Supplement 1***).

To elucidate how crowding promotes Dz activity, we first focused on the physiochemical properties density, viscosity, pH, and the theoretical, average distance between cosolutes *D* (***Figure 6***, ***Figure 6***—***figure Supplement 1***). Since none of the crowded solutions presented significant changes in pH and density, we suggest that viscosity-related alterations contribute in elevating Dz activity. Here, ‘optima’ provide similar solution viscosities for one cofactor (***Figure 6, Figure 3***, ***Figure 3***— ***figure Supplement 1***). In this regard, size and concentration of the respective cosolute not only affects the solution’s viscosity, but also determines the average distance between two cosolute molecules (***Figure 6***—***figure Supplement 1***), thus the volume available to the Dz:RNA complex. This makes structural information necessary to better understand steric influence by volume exclusion, which we addressed using FCS (***Figure 7***) and SAXS (***Figure 8***). From our FCS data we observe that a *r*_h, TDz_ ~ 1.81 − 2.0 nm in combination with *D* of 6.50 nm is associated with activity favoring conditions for Mg^2+^-induced catalysis (***Figure 7A, Figure 7B***). In presence of NaCl, preferences in *D* extend to 8.41 nm. For Mn^2+^-induced RNA cleavage, promoting conditions are associated with *D* ~ 8.40 − 11.1 nm and *r*_h, TDz_ ≳ 2.0 nm (***Figure 7***—***figure Supplement 2***). Under conditions with NaCl, an ‘optimum’ cavity size seems to be 11.14 nm. In general, conditions that cause the Dz:RNA complex to compress to a *r*_h, TDz_ below 1 nm tend to restrict activity. This accounts for high concentrations of larger PEG molecules (0.18 g/mL PEG 20000, ***Figure 7E***), which we assign to a higher degree of volume exclusion, thus an increased potential of steric repulsion (***Dupuis et al., 2014; Kilburn et al., 2010; Sugimoto, 2014***). Although we observe that certain combinations of viscosity and *D* can create a favoring surrounding, our data do not indicate a correlation between physiochemical properties and the corresponding *r*_h_ of the Dz:RNA complex. This makes us believe that other environmental factors, for instance osmotic pressure (***DasGupta, 2020***) or dielectric properties (***Fiorini et al., 2015; Nasir and Al Ahmad, 2020***), also impact the DNAzyme, which should be evaluated. In addition, since we see that one *r*_h, TDz_ can account for different degrees in Dz activity, we propose that Dz:RNA complex structure is impacted less by molecular crowding. This is also supported by the SAXS data, which indicated only small deviations in the Dz:RNA complex in presence of crowded conditions (***Figure 8***). Besides, the results show that interparticle interactions for the Dz:RNA complex change in presence of MgCl_2_. The published data by ***Borggräfe et al. (2021***) revealed that after binding of monovalent ions to the complex, its saturation with Mg^2+^ is required to achieve structural arrangements that result in the active conformation. Taken both observations into account, this raises the question how the complex behaves structurally under crowded conditions in a cofactor-bound state, as FCS measurements were performed in absence of Mg^2+^ and Mn^2+^.

For the first time we show that also the activity of the Dz is impacted in presence of a more cell-like surrounding, which is indicated to be dependent on the cosolute’s properties and correlates with a previous study on the 8-17 DNAzyme variant 17E by ***Nakano et al. (2017***). In addition, we provide first information on structural changes of the Dz:RNA complex under crowded conditions and highlight that cosolute concentration is one critical factor for Dz activity. To the best of our knowledge, this has not been shown for the Dz in presence of the selected cosolutes yet. The mentioned study by ***Nakano et al. (2017***) supports the significance of cofactor concentration. By testing 10, 20 and 30 % of PEG 200, PEG 8000, and EtOH on DNAzyme activity and stability, they were able to observe that the extent, to which both are altered depends on the cosolute concentration. Here, catalytic activity and DNA stability presented a high level of correlation (***Nakano et al., 2017***). In this study, we point out that few crowded conditions are able to promote Dz activity that seem to achieve a catalytically relevant degree of structural adaption of the Dz:RNA complex, although indicated as minor according to the FCS and SAXS data. If connected to promoting conditions in Mg^2+^- or Mn^2+^-induced cleavage, *r*_h, TDZ_ tendencies slightly differ between the two metal ions. Since both ions presented different interplays with monovalent ions during catalysis (***Rosenbach et al., 2021***), this raises the question of cofactor-dependent conformational differences between Dz:RNA complexes that would explain variations in the preference of crowded conditions. Based on our data, we refer to observations by ***Paudel et al. (2018***) and also propose that the cavity size mediated by crowding influences Dz efficiency. But we have to note, that *D* calculations include a correction factor, whose underlying concept needs to be reviewed for the rest of the PEG variants in this study. Since cavity size distribution here is homogeneous, but is most likely heterogeneous, given the molecular diversity in cells, this further questions whether the impact on Dz activity changes in a heterogeneous environment. Assays in this study were performed with pre-formed complexes under single-turnover. Although we suggest that the above mentioned alterations in complex size and solution properties will negatively impact complex association and dissociation under multiple turnover, validation is required. Our FCS data did not show notable differences between T and TDz under crowded conditions (***Figure 7***, ***Figure 7***—***figure Supplement 1***). At this point, this leaves us to speculate that RNA substrate and Dz:RNA complex are affected to a similar extent, despite size differences and estimated volume occupancy. Since FCS analyses have been performed based on the fluorescence of the single-labeled RNA substrate, further investigations are required to exclude multiple species in TDz samples. Despite first structural insights in this study, high resolution structural data is mandatory that would provide important insights into the reaction mechanism and contribute to a better understanding of conformational changes within the Dz:RNA complex that result from interactions with the respective cosolute. In this regard, possible rearrangements of catalytically relevant residues within the catalytic core are of great interest. Moreover, detailed structural data will also help understand the observed differences between Mg^2+^- and Mn^2+^-based approaches.

## Materials and methods

### Oligonucleotide sequences

DNA and RNA oligonucleotides were commercially obtained from biomers.net (Ulm, Germany). Single-labeled RNA substrates used in the stability assay followed by denaturing urea-PAGE contained a 6-carboxyfluorescein (6-FAM) molecule at the 5’ end. Single-labeled RNA substrates used for fluorescence correlation spectroscopy (FCS) measurements contained an Atto647N molecule at the 5’ end. RNA substrates in Förster resonance energy transfer (FRET)-based activity assays were double-labeled with a 6-FAM molecule at the 5’ end and the Black Hole Quencher-1 (BHQ-1) at the 3’ end. In terms of FCS and small angle x-ray scattering (SAXS) studies, the RNA substrate was additionally stabilized by substituting the 2’-OH group of the ribose moiety of the guanosine at the cleavage site with a fluorine atom (2’F) to avoid cleavage (***Borggräfe et al., 2021; Rosenbach et al., 2021***). The Dz used in this study was designed to specifically bind the human prion protein (PrP) mRNA and perform cleavage at position 839 (***Victor et al., 2018***). The sequences for Dz and RNA substrates are listed in ***Table 1***.

### Cosolutes to simulate crowded conditions

Cosolutes to simulate crowded conditions *in vitro* were obtained commercially and used at concentrations of 0.03, 0.07, 0.11, 0.15 and 0.18 g/mL, respectively. The concentration range was selected according to published studies (***Fiorini et al., 2015; Paudel et al., 2018***) and estimated total concentrations of biomolecules in cells (50-400 g/L; ***Zimmerman and Minton, 1993***). Samples containing osmolytes were prepared either with betaine (BioUltra, ≥ 99 %, Sigma-Aldrich, St. Louis, MO, USA) or ectoine (≥ 95 %, HPLC, Sigma-Aldrich, St. Louis, MO, USA). In addition, we used ethylene glycol (EG, ≥ 99 %, Muskegon, MI, USA) and different variants of polyethylene glycol (PEG) with average molecular masses (*M*_W, av_) of 200 (PEG 200, for synthesis), 600 (PEG 600, for synthesis), 1000 (PEG 1000, for synthesis), 3000 (PEG 3000, BioUltra), 3350 (PEG 3350, BioUltra), 4000 (PEG 4000, for synthesis), 6000 (PEG 6000, BioUltra), 8000 (PEG 8000, BioUltra) and 20000 (PEG 20000, (stabilized) for synthesis). PEG 200 to 8000 were obtained from Sigma-Aldrich (St. Louis, MO, USA) and PEG 20000 from Merck (Darmstadt, Germany). In terms of polysaccharides, dextran 5000 (dex 5000,*M*_W, av_ = 5 kDa) and 12000 (dex 12000, *M*_W, av_ = 12 kDa, analytical standard for GPC, Sigma-Aldrich, St. Louis, MO, USA) were used. Stocks were prepared as 50 % (g/ml) solutions in either water (18.2 MΩ) or 50 mM Tris-HCl, pH 7.5.

### Stability assay with single-labeled RNA substrates followed by denaturing urea-PAGE

Stability assays were performed with 0.4 µM 6-FAM-labeled RNA substrate and 0.4 µM Dz (single-turnover) in 50 mM Tris-HCl, pH 7.5 in presence and absence of 100 mM sodium chloride (NaCl) and cosolute. Prior to cleavage, RNA substrate and Dz were denatured in buffer without Mg^2+^ at 73 °C for 5 min, followed by an incubation of 10 min at room temperature for renaturation. RNA cleavage was induced with 1 mM magnesium chloride (MgCl_2_) and incubated for 3 h at 37 °C. Samples taken for further analysis were quenched in 94 % formamide, 25 mM ethylenediamine tetraacetic acid (EDTA), pH 8.0, 0.02 % Bromphenol blue (w/v), and 0.02 % Xylene cyanol (w/v) and heated at 95 °C for 10 min. Sample separation was carried out using 18 % polyacrylamide gels with 7 M urea buffered with Tris-borate EDTA buffer (TBE) for 45 min at 20 W (***Kirchgässler et al., 2022***). Prior to sample separation by gel electrophoresis, gels were preheated for 30 min at 20 W to avoid temperature inhomogeneities. 6-FAM labeled RNA substrates were detected and visualized by fluorescence, which allows detection of uncleaved RNA substrates and one of two cleavage products that contains the 6-FAM molecule at the 5’ end (5’-fragment). In case of unlabeled RNA substrates, visualization was provided by staining in TBE buffer that contains a dilution of 1:30,000 of GelRed nucleic acid gel stain (10,000X in H_2_O, Biotium, Fremont, CA, USA) for 1 h at room temperature. Images were acquired using the ChemiDoc MP System (Bio-Rad, Hercules, CA, USA).

### FRET-based cleavage assays

Cleavage assays were carried out using a double-labeled RNA substrate (T^FRET^, Table 1). Metal ion induced RNA cleavage allows detection of an increase in fluorescence over time. Assays were performed with final concentrations of 0.1 µM RNA substrate and 0.1 µM Dz (single-turnover) in 50 mM Tris-HCl, pH 7.5 and 0.1 mM EDTA with different concentrations of cofactors (MgCl_2_ : 0.5, 3 and 5 mM; manganese(II) chloride (MnCl_2_) : 0.5 mM) at 37 °C. Measurements were carried out in presence and absence of 100 mM NaCl and cosolute.

0.8 µM RNA substrate and Dz were denatured in buffer for 5 min at 73 °C, followed by an incubation of 10 min at room temperature to allow renaturation. Samples were prepared at higher concentrations to facilitate complex formation. Initial denaturation was performed in either presence or absence of NaCl, but absence of cosolute and MgCl_2_ or MnCl_2_. During renaturation, different cosolute concentrations were added to the wells of a 384 well non-binding microplate (Greiner Bio-One, Kremsmünster, Austria) in buffer and a total volume of 15 µL. Nucleic acid stocks were diluted with buffer to a construct concentration of 0.2 µM. Next, 20 µL of solution were added to the wells. Each measurement included a control that contained the RNA substrate only in absence of cosolute. The plate was sealed with tape (Polyolefin Acrylate, Thermo Scientific, Waltham, MA, USA), placed inside the plate reader (CLARIOStar, BMG Labtech, Ortenberg, Germany), and equilibrated to 37 °C for 30 min. Prior to measurement, the sealing tape was removed and RNA cleavage was induced by automatic injection of 5 µL of cofactor stock solution (MgCl_2_ or MnCl_2_) after 15 cycles of measurement. Data points were acquired with cycle times of either 8 s (0.05 mM MgCl_2_), 7 s (0.1 mM MgCl_2_), 6 s (0.2 mM MgCl_2_), 5 s (0.3 mM MgCl_2_), 4 s (0.5 mM MgCl_2_ and 0.01 mM MnCl_2_), 3 s (0.05 mM MnCl_2_), 2 s (3 and 5 mM MgCl_2_, 0.1, 0.2 and 0.25 mM MnCl_2_), or 1 s (0.3, 0.4 and 0.5 mM MnCl_2_) depending on the speed of reaction. Assays in competition with ATP were carried out with 0, 0.2, 0.5, 0.8, 1, 1.5 and 2 mM ATP at 0.5 mM MgCl_2_ in 50 mM Tris-HCl, pH 7.5, 0.1 mM EDTA, and 100 mM NaCl. Measurements were performed with 1000 cycles and excitation and emission wavelength were set to *λ*_ex_ = 484 nm and *λ*_ex_ = 530 nm.

Experimental data was fitted and plotted using graphics layout engine (GLE, https://glx.sourceforge.io/). Fluorescence intensity curves were fitted using

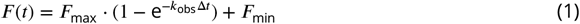

with *F*_max_ and *F*_min_ as the maximum and minimum fluorescence in curves, *k*_obs_as the observed rate constant, and Δ*t* = *t* − *t*_injection_ as time difference after subtraction of RNA only fluorescence and data correction for a time-dependent linear decrease in fluorescence, most likely due to photo-bleaching.

To evaluate the impact of molecular crowding on Dz activity, the relative rate constant *k*_rel,*i*_ was calculated

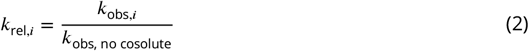

### FCS for diffusion analysis

Sample preparation for FCS analyses was carried out in sets of RNA substrate only (T) and Dz:RNA complex (TDz) for each tested condition using the stabilized RNA substrate T^Atto647N^ (Table 1). In absence of cosolute, stocks containing 400 nM of T^Atto647N^ and Dz^WT^ in 50 mM Tris-HCl, pH 7.5 and 0.1 mM EDTA were incubated for 5 min at 73 °C for denaturation, followed by an incubation of 10 min at room temperature to facilitate complex formation. Subsequently, stocks of T and TDz were diluted 1:10 with 50 mM Tris-HCl, pH 7.5 and 0.1 mM EDTA. Samples were prepared separately in reaction tubes with total volumes of 210 µL. Here, cosolutes and buffer were added first. 10.5 µL of diluted stock of T or TDz were added immediately before measurement to reach a final concentration of 2 nM. Samples were mixed thoroughly and 200 µL each were transferred into wells of sterile NuncTM Lab-TekTM II chambered coverglass systems (8 chambers, Nunc, Thermo Scientific, Rochester, NY, USA). Closed sample chambers were incubated for 20 min at 37 °C prior to measurement. The following conditions have been analyzed and were selected presenting all three types of cosolutes, different degrees of volume exclusion, and Dz activity: 50 mM Tris-HCl, pH 7.5 and 0.1 mM EDTA (buffer only), 0.03 g/mL PEG 4000, 0.07 g/mL PEG 3000 and 3350, 0.11 g/mL PEG 8000, 0.15 g/mL PEG 200, 600, 3000, and 4000, and 0.18 g/mL betaine, ectoine, dextran 12000, EG, and PEG 20000.

FCS data were collected using an Olympus FLUOVIEW FV3000 (Olympus Europa SE & Co. KG, Hamburg, Germany) confocal laser scanning microscope with a high numerical aperture objective (UPLSAPO60XW/1.2). The microscope is equipped with a multichannel picosecond event timer (HydraHarp 400, PicoQuant) for single-point time correlated single photon counting (TCSPC) measurements. Samples were excited using a 640 nm CW laser (OBIS, Coherent) and the fluorescence signal was split for its polarization using a polarizing beamsplitter. The parallel and perpendicular signal was detected using two identical bandpass filters (690/70, AHF Analysentechnik, Germany) in front of two avalanche photodiodes (SPCM-AQRH-14-TR module, Excelitas Technologies) to enable detector cross-correlation and after pulse correction. FCS measurements were performed at 37 °C and with a laser intensity of 16.7 µW. All conditions (see above) were measured in triplicates (*n* = 3) to check for reproducibility, of which each measured triplicate consists of five measurements. From data of the obtained time traces, cross-correlation curves were calculated using the SymPhoTime 64 software (PicoQuant, Berlin, Germany). The cross-correlation curve of five independent measurements was averaged and fitted. The fitting model *G*(*τ*) accounts for the free diffusion of the molecule of interest within and the signal fluctuations due to the population of a potential triplet state of the used fluorophore (***Lakowicz, 2006; Yu et al., 2021***) according to

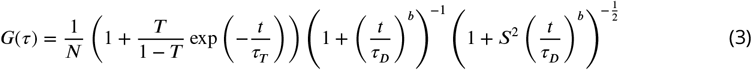

with *N* as average number of molecules within the confocal volume, *T* as triplet fraction, *τ*_*D*_ as diffusion time, *τ*_*T*_ as characteristic correlation time of the triplet state, *b* as anomalous or subdiffusion exponent due to molecular crowding (***Höfling and Franosch, 2013***), and *S* = 0.2 as structure factor, i. e., the aspect ratio of the 3D gaussian approximation of the confocal volume. The triplet fraction accounts for the varying population of the triplet state of the fluorophore used. We calibrated the confocal volume using a dye with known diffusion coefficient *D* and calculated the focus radius 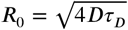 using the diffusion coefficient of Atto655 with *D*_37 °C_ = 5.62 ⋅ 10^−10^ m^2^s^−1^ (***Müller et al., 2008***) due to structural similarity between Atto655 and Atto647. We determined *b* = 1 with calibration measurements of the free dye, as expected. In the presence of cosolutes, however, we found *b* = 0.9, which we attribute to subdiffusion of the nucleic acid molecules in the crowded solution. The offset of the autocorrelation curve has been found to be zero. The Stokes radius, thus, hydrodynamic radius *r*_*h*_ of the molecule of interest was calculated according to Stokes-Einstein

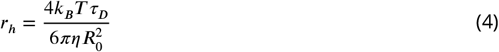

with *k*_*B*_ as Boltzmann constant, *T* the absolute temperature and *η*(*T*) as the experimentally determined dynamic viscosity of the crowded solution.

### Calculation of *r*_**h**_, *r*_**g**_, **and cavity size** *D*

For determining the size of PEG particles, the radius of gyration *r*_g_ was calculated based on light scattering results (***Devanand and Selser, 1991***) following the equation

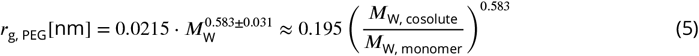

with *M*_W, monomer_ ≈ 44 g/mol and considering volume exclusion effects, thus no overlap of cosolutes. The theoretical relation between the according hydrodynamic radius *r*_h_ and *r*_g_ for a linear polymer in a good solvent is *r*_h_/*r*_g_ = 0.64 (***Linegar et al., 2010; Teraoka, 2002***). *r*_h_ for PEG particles was calculated based on ***Devanand and Selser (1991***)

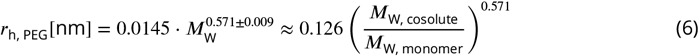

Hydrodynamic radii *r*_h_ for dextran molecules were calculated based on previous experiments by ***Hernández et al. (2017***)

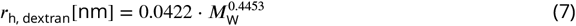

with *M*_W_ as the average molecular weight of the cosolute particle.

The theoretical, average distance between cosolutes (cavity size, *D*) was calculated based on the assumption of a simplified hard sphere model, (***Kang et al., 2015***)

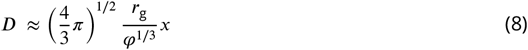

where *φ* is the volume fraction of all cosolute particles in solution, which depends on size of the particle, its number in solution, and its mass concentration.

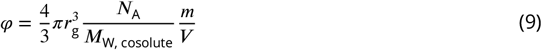

The correction factor *x* is introduced to adjust differences between *r*_g_ and *r*_h_, which was determined empirically for cosolutes EG (5.8), PEG 1000 (2.8), PEG 8000 (1.3), and PEG 35000 (0.65) in a previous ribozyme-based approach (unpublished). Theoretical correction factors for the remaining PEG variants in this study have been calculated according to the fit model

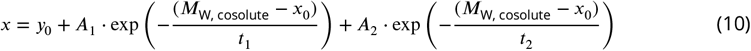

with *y*_0_ = 0.65851, *A*_1_ = 2.0757, *x*_0_ = 62, *t*_1_ = 6999.927, *A*_2_ = 3.0757, and *t*_2_ = 387.688.

### Structural analysis by SAXS

Small angle X-ray Scattering (SAXS) measurements were performed with final concentrations of 5 g/mL of Dz:RNA complex. In absence of cosolute and MgCl_2_, Dz and RNA substrate (T^2’F^) were denatured in 50 mM Tris-HCl, pH 7.5, 0.1 mM EDTA, and 100 mM NaCl at 73 °C for 5 min, followed by an incubation of 10 min at room temperature. Next, cosolutes or MgCl_2_ dissolved in 50 mM Tris-HCl, pH 7.5, 0.1 mM EDTA, and 100 mM NaCl were added to reach the final complex concentration for measurement.

SAXS experiments were performed at the Jülich Center for Neutron Science (JCNS) at Forschungszentrum Jülich, Germany. The instrument “Ganesha-Air” from SAXSLAB/XENOCS was used. The X-ray source is a D2-MetalJet (Excillum) with Ga–K*α* radiation (wavelength *λ* = 0.134 14 nm). The data were acquired with a position-sensitive detector (PILATUS 300 K, Dectris). After calibration with silver behenate, the distance from the sample to the detector was set to cover a *Q*-range of 0.1 − 6 nm^−1^. Temperature was set to 20 °C.

The radius of gyration *r*_g_ was determined by a fit to the measured intensity

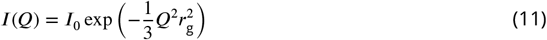

for *Qr*_g_ using the Guinier model (***Feigin and Svergun, 1987***) with magnitude of the scattering vector

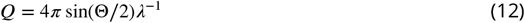

and scattering angle Θ.

Ellipsoidal scattering formfactor *P* (*Q*) for a sphere with semiaxes *r*_a_ = *eR* (axis of revolution) and *r*_b_ = *R, e* = *r*_a_/*r*_b_ and contrast Δ*ρ* to the solvent (***Pedersen, 1997***) is

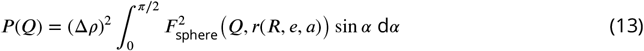

with

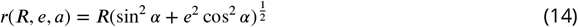

and

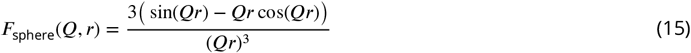

The double Yukawa potential of a particle with radius *r* at a position *r*′ is

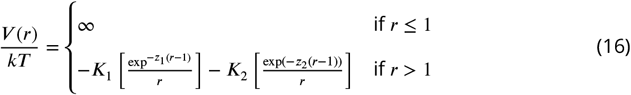

with *z*_*i*_ as inverse screening length and potential *K*_*i*_ at the surface. For *K* > 0 we have attraction while *K* < 0 means repulsion. ***Liu et al***. described the corresponding structure factor *S*(*Q*) within the MSA closure (***Liu et al., 2005***). Data analysis was done using the Python-based tool Jscatter (***Biehl, 2019***).

### Analysis of physiochemical solution properties

All physiochemical solution properties in presence and absence of cosolutes were determined in 50 mM Tris-HCl and pH 7.5, 0.1 mM EDTA with samples prepared in triplicates (*n* = 3) and at 37 °C, respectively. Density measurements have been performed using a DM450M densiometer (Anton-Paar, Ostfildern-Scharnhausen, Germany). Determining the dynamic and kinematic viscosity of solutions was carried out using a rolling ball viscosimeter Lovis 2000 M (Anton-Paar, Ostfildern-Scharnhausen, Germany) using a glass capillary (short) with a diameter of 1.59 mm (un-calibrated) and a steel ball of 1.5 mm diameter in size and 7.69 g/cm^3^ in density (Mat no 73109, steel: 1.4125, Anton-Paar, Ostfildern-Scharnhausen, Germany). Prior to measurement, calibration was performed with water (18.2 MΩ) at 20 °C and a maximal variation coefficient of 0.1 %. In terms of pH analyses, samples were incubated for 4 h at 37 °C prior to measurement. pH values were determined using a standard pH meter (FiveEasy, Mettler-Toledo, Gießen, Germany) and glass combination electrode (REF 238140, BioTrode, Hamilton, Bonaduz, Switzerland).

## Supporting information

### Supplementary figures

- Figure 2—figure supplement 1. Urea-PAGE of samples after stability analysis in presence of 0.18 g/ml of the remaining cosolutes
- Figure 3—figure supplement 1. Selection of cosolute concentrations that included conditions with promoting properties on Dz activity at 0.5 mM MgCl_2_
- Figure 3—figure supplement 2. Effects of crowded conditions on Dz activity in presence and absence of 100 mM NaCl at 0.5 mM MnCl_2_
- Figure 3—figure supplement 3. Dz activity depending on MgCl_2_ concentration
- Figure 4—figure supplement 1. Dz activity in presence of crowding and competition with ATP at 0.5 mM MgCl_2_ and 100 mM NaCl
- Figure 6—figure supplement 1. Hydrodynamic radius (*r*_h_, **A**), radius of gyration (*r*_g_, **B**), and theoretical, average distance between cosolutes (*D*, **C**) depending on *M*_W_ and concentration
- Figure 7—figure supplement 1. Diffusion time *τ*_D_ of RNA substrate (T) and Dz:RNA complex (TDz) in absence and presence of cosolutes determined by FCS.
- Figure 7—figure supplement 2. Comparison of the Dz:RNA complex’s hydrodynamic radius *r*_h_, *D*, and relative Dz activity for Mn^2+^-induced RNA cleavage.

### Supplementary table

- Figure 6—table supplement 1. Density *ρ* and dynamic viscosity *η* at 37 °C of additional samples used in FCS measurements with cosolute concentrations less than 0.18 g/ml
- Figure 6—table supplement 2. *D* of all used PEG variants at a concentration range of 0.03 to 0.18 g/mL

## Acknowledgments

We thank apl. Prof. Dr. M. Schmitt from the Institute for Physical Chemistry I at Heinrich-Heine-Universität Düsseldorf for providing access to the densiometer to perform density analyses. We would also like to acknowledge the Center for Advanced Imaging (CAi) at Heinrich-Heine-Universität Düsseldorf for providing access to the facilities’ microscope and especially apl. Prof. Dr. S. Weidtkamp-Peters for the assist and support with setting up the FCS experiments. Funding for instrumentation: Olympus FV3000 Confocal Laserscanning Microscope: DFG-INST 1358/44-1 FUGB.

---

**Figure 2—figure supplement 1.**
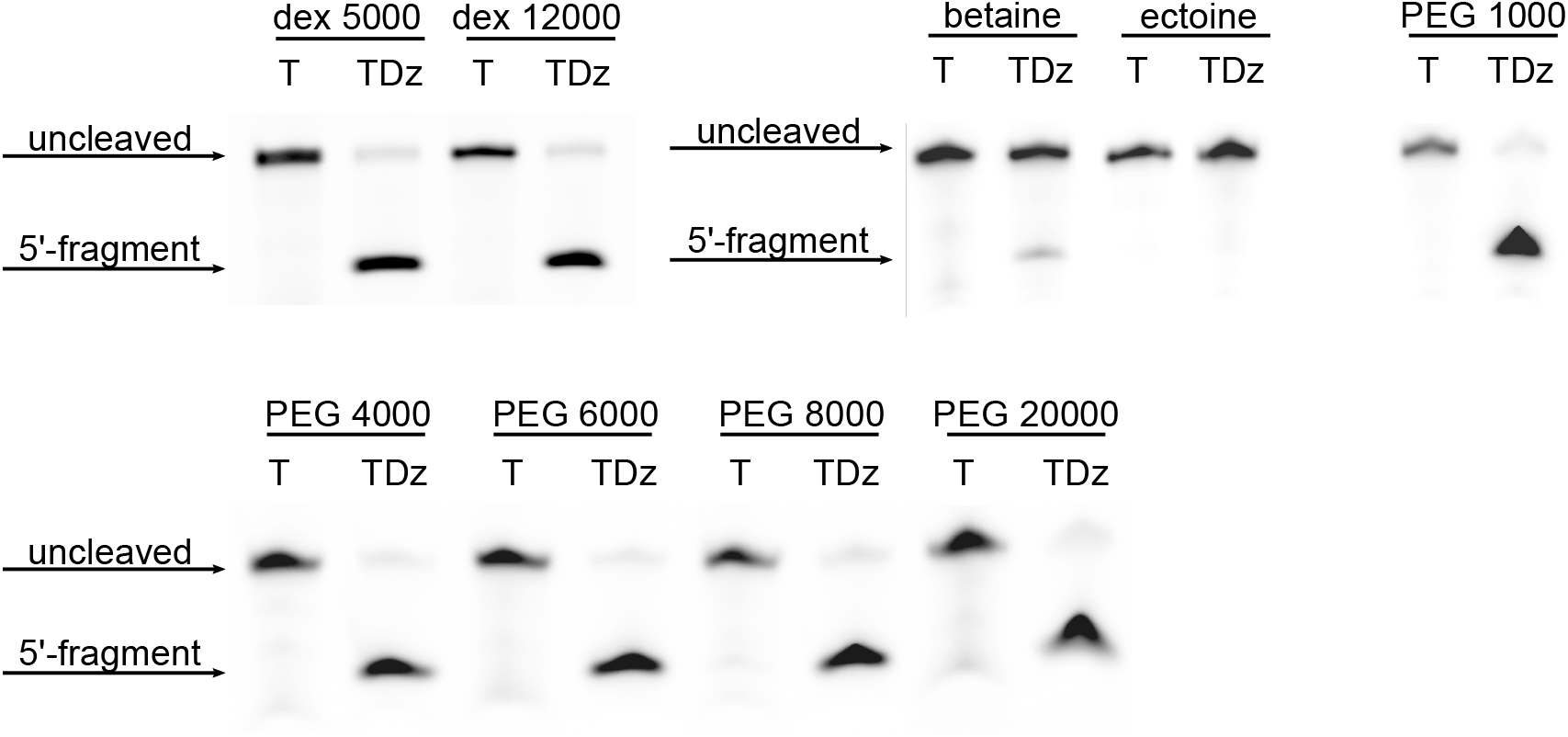
Urea-PAGE of samples after stability analysis in presence of 0.18 g/ml of the remaining cosolutes. Conditions were analyzed in pairs of equally treated samples containing the RNA substrate only (T) and the DNAzyme:RNA complex (TDz). Assays were performed using the T^FAM^ and Dz^WT^ constructs (***Table 1***). Fluorescence detection and visualization of 6-FAM labeled RNA substrates and the 5’ cleavage fragment.

**Figure 3—figure supplement 1.**
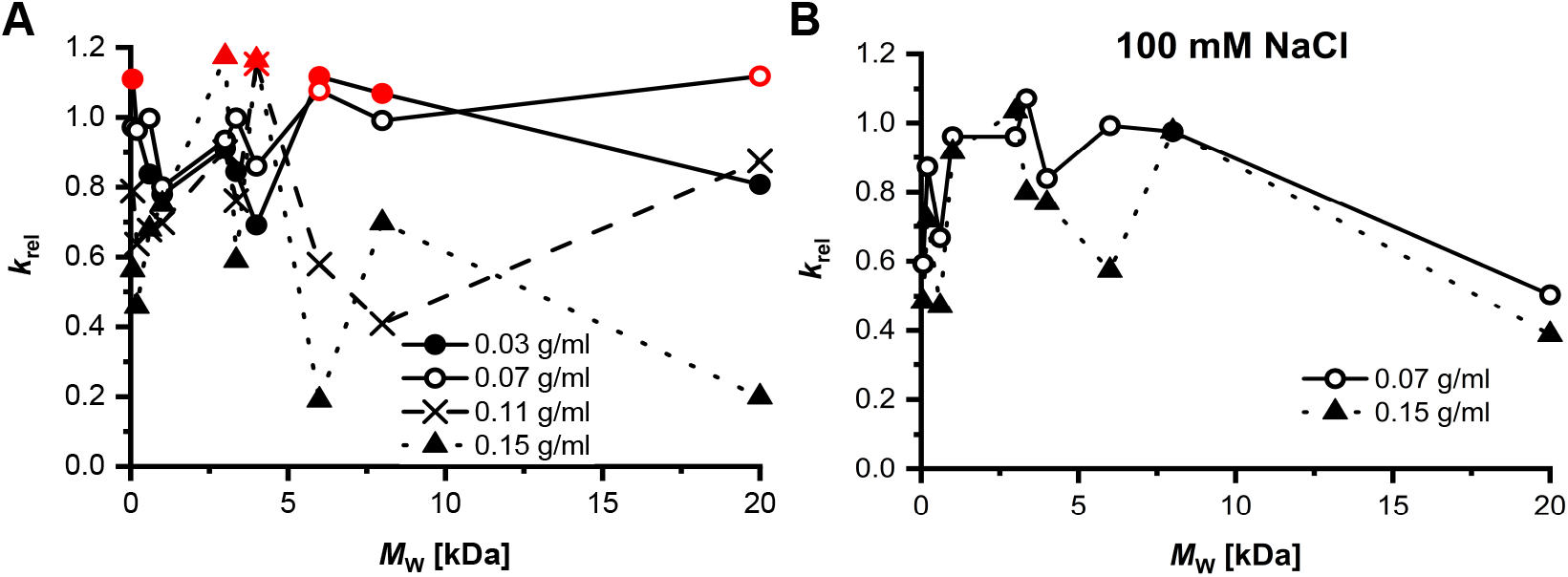
Selection of cosolute concentrations that included conditions with promoting properties on Dz activity at 0.5 mM MgCl_2_. **A** *k*_rel_ of respective concentrations of PEG variants in absence of NaCl. Promoting conditions with *k*_rel_ ≥ 1.10 are marked in red (•). **B** *k*_rel_ for 0.07 and 0.15 g/mL of PEG variants in presence of 100 mM NaCl. Promoting conditions include 0.07 g/ml of PEG 3350 (*k*_rel_ = 1.07) and 0.15 g/ml of PEG 3000 (*k*_rel_ = 1.03).

**Figure 3—figure supplement 2.**
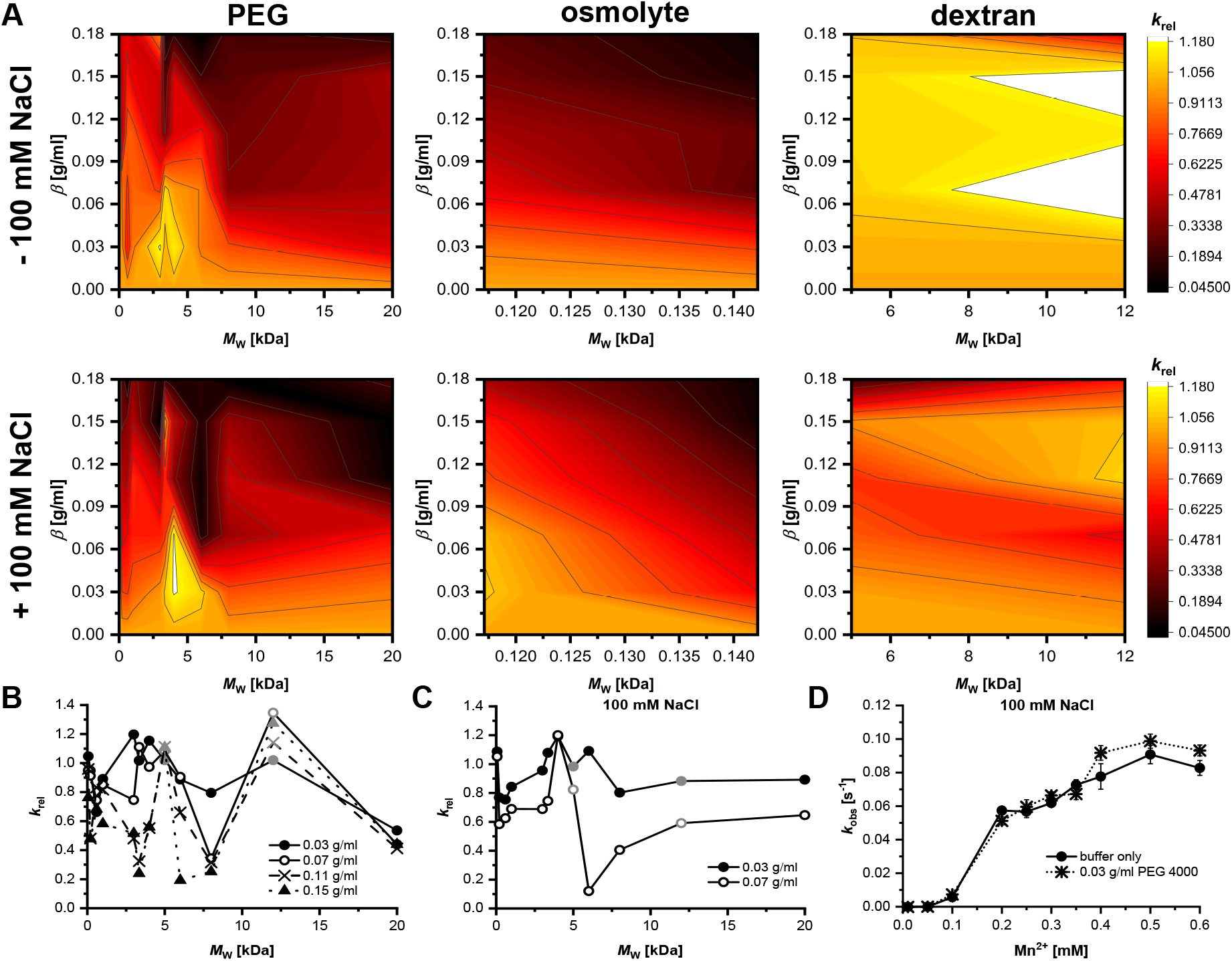
Effects of crowded conditions on Dz activity in presence and absence of 100 mM NaCl at 0.5 mM MnCl_2_. Dz activity was determined in FRET-based activity assays. **A** Relative activity in presence of either PEG, osmolytes betaine and ectoine or dextran. Relative rate constants *k*_rel_ (Equation 2) are based on determined rate constants *k*_obs,*n*=14_ = (0.1050 ± 0.0299) s^−1^ and *k*_obs,*n*=14_ = (0.0459 ± 0.0076) s^−1^ with and without 100 mM NaCl, respectively. **B** *k*_rel_ for different concentrations of PEG and dextran in absence of NaCl that include promoting conditions on Dz activity. Samples containing dextran are highlighted in grey (•). Conditions that enhanced Dz activity (*k*_rel_ > 1.10) include: 0.03 g/mL PEG 3000, PEG 3350, and PEG 4000; 0.07 g/mL PEG 3350 and dextran 12000; 0.11 and 0.15 g/mL dextran 5000 and 12000. **C** *k*_rel_ for different concentration of PEG and dextran variants in presence of 100 mM NaCl. Samples containing dextran are highlighted in grey (•). Conditions that that enhance Dz activity (*k*_rel_ > 1.10) include: 0.03 and 0.07 g/mL PEG 4000. **D** Dz activity as *k*_obs_[s^−1^] in absence of crowder (buffer only) and presence of 0.03 g/mL of PEG 4000 (most promoting condition, *k*_rel_ = 1.13) at MnCl_2_ concentrations 0.01 − 0.6 mM and 100 mM NaCl.

**Figure 3—figure supplement 3.**
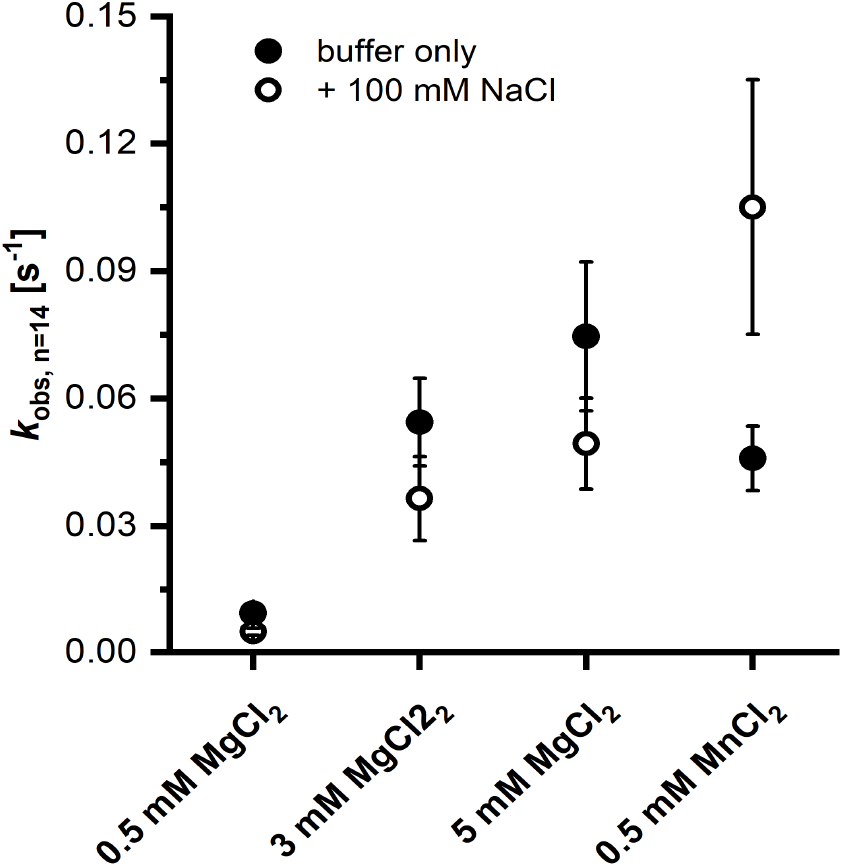
Dz activity depending on MgCl_2_ concentration. Mean values of *k*_obs_[s^−1^] for Dz activity in buffer only with and without 100 mM NaCl. Mean values were calculated based on *n* = 14 samples, respectively.

**Figure 4—figure supplement 1.**
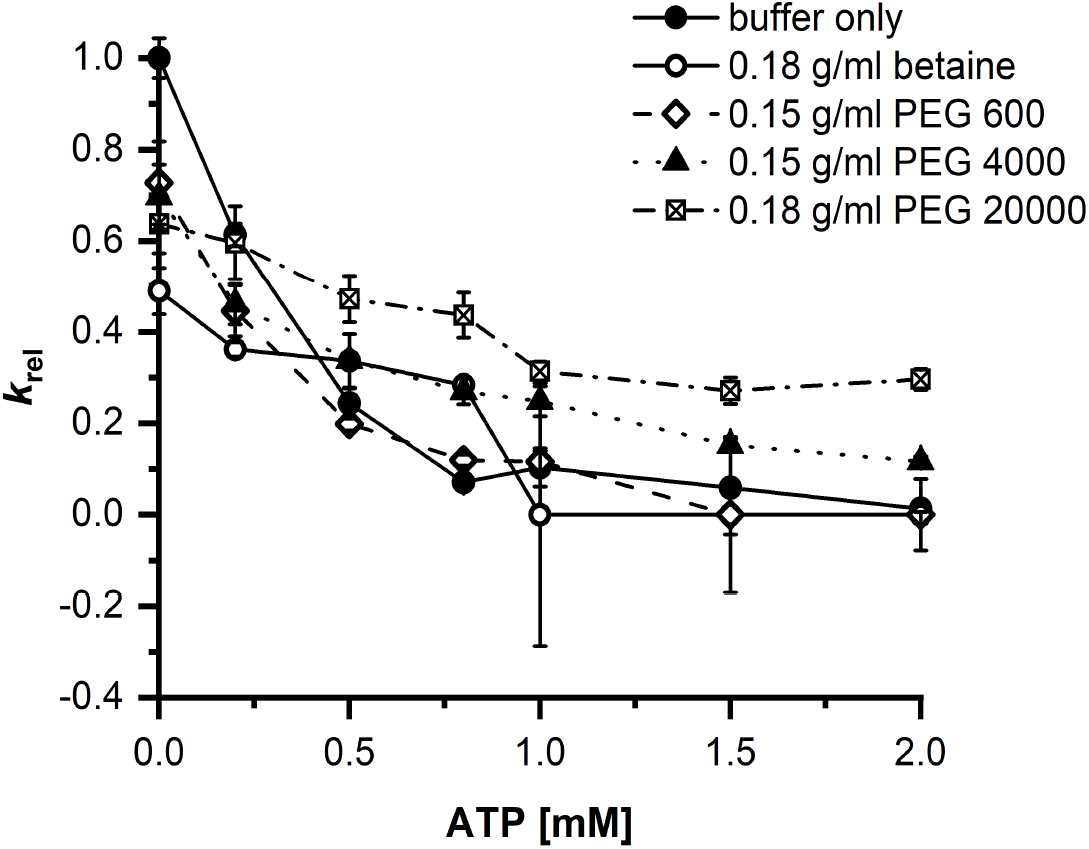
Dz activity in presence of crowding and competition with ATP at 0.5 mM MgCl_2_ and 100 mM NaCl. Rate constants (*k*_obs_) were put into relation to the activity in absence of cosolute and ATP (*k*_rel_, (2)) with 100 mM NaCl (*k*_obs,*n*=3_ = (0.0035 ± 0.0001) s^−1^).

**Figure 6—figure supplement 1.**
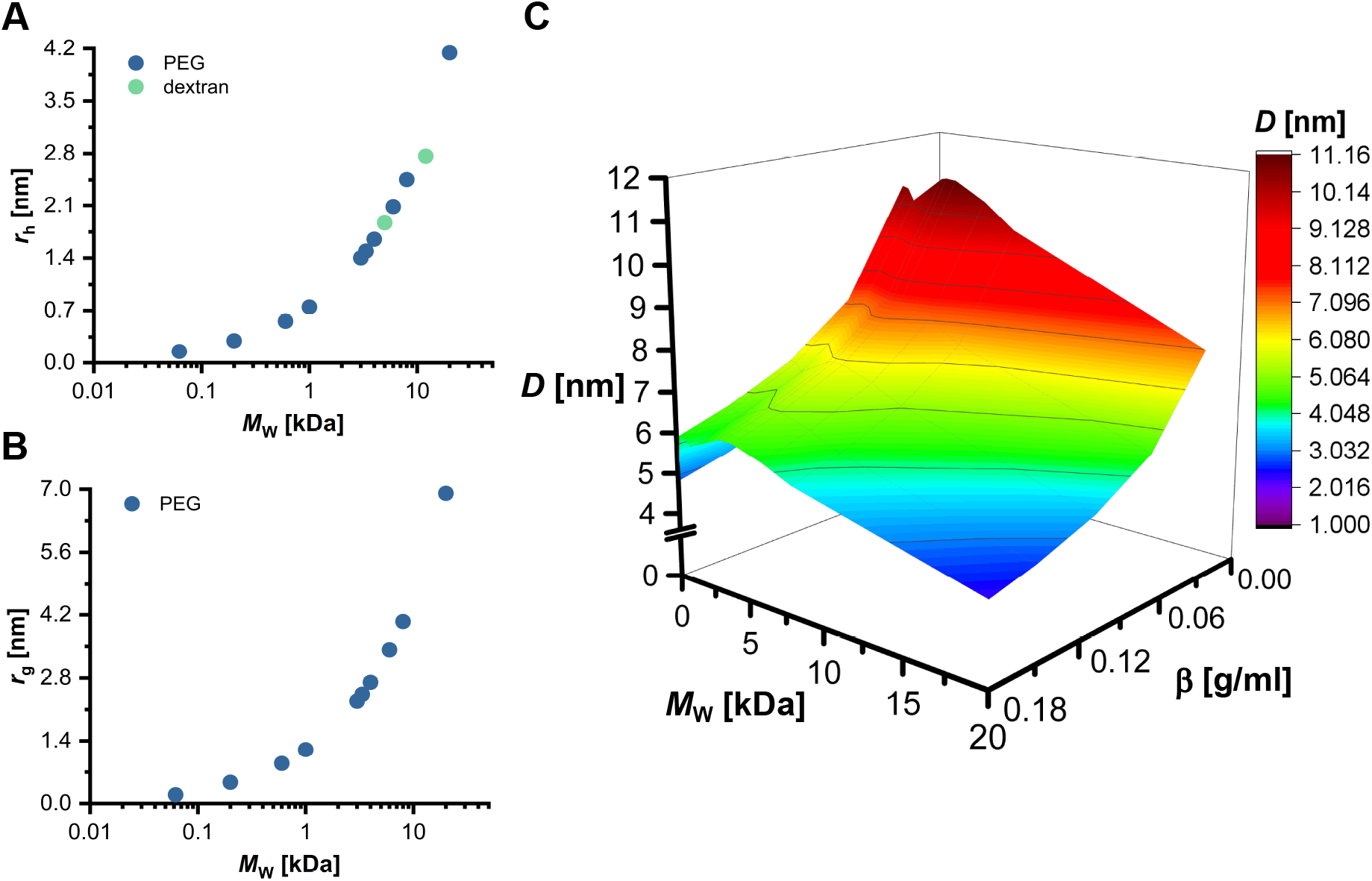
Hydrodynamic radius (*r*_h_, **A**), radius of gyration (*r*_g_, **B**), and theoretical, average distance between cosolutes (*D*, **C**) depending on *M*_W_ and concentration. **A** *r*_h_ were calculated based on ***Equation 6*** and ***Equation 7*. B** *r*_g_ of PEG were calculated based on ***Equation 5*. C** *D* between PEG molecules in solution was calculated based on the *r*_g_ and under the assumption of a hard sphere model (***Equation 9, Equation 8***, and ***Equation 10***). *D* in buffer only is assumed to be 100 nm.

**Figure 6—table supplement 1.** Density *ρ* and dynamic viscosity *η* at 37 °C of additional samples used in FCS measurements with cosolute concentrations less than 0.18 g/ml.

| Sample | $\rho_{37^\circ\text{C}}$ [g/cm<sup>3</sup>] | $\eta_{37^\circ\text{C}}$ [mPas] |
| --- | --- | --- |
| PEG 200, 0.15 g/mL | 1.0181 ± 0.000 03 | 1.1553 ± 0.002 62 |
| PEG 600, 0.15 g/mL | 1.0199 ± 0.000 75 | 1.4672 ± 0.084 75 |
| PEG 3000, 0.07 g/mL | 1.0061 ± 0.000 02 | 1.4940 ± 0.005 35 |
| PEG 3000, 0.15 g/mL | 1.0191 ± 0.000 08 | 3.2453 ± 0.000 94 |
| PEG 3350, 0.07 g/mL | 1.0063 ± 0.000 05 | 1.4797 ± 0.016 74 |
| PEG 4000, 0.03 g/mL | 1.0004 ± 0.000 16 | 1.0320 ± 0.002 16 |
| PEG 4000, 0.15 g/mL | 1.0197 ± 0.000 17 | 3.5538 ± 0.267 21 |
| PEG 8000, 0.11 g/mL | 1.0125 ± 0.000 58 | 3.7220 ± 0.056 45 |

**Figure 6—table supplement 2.**
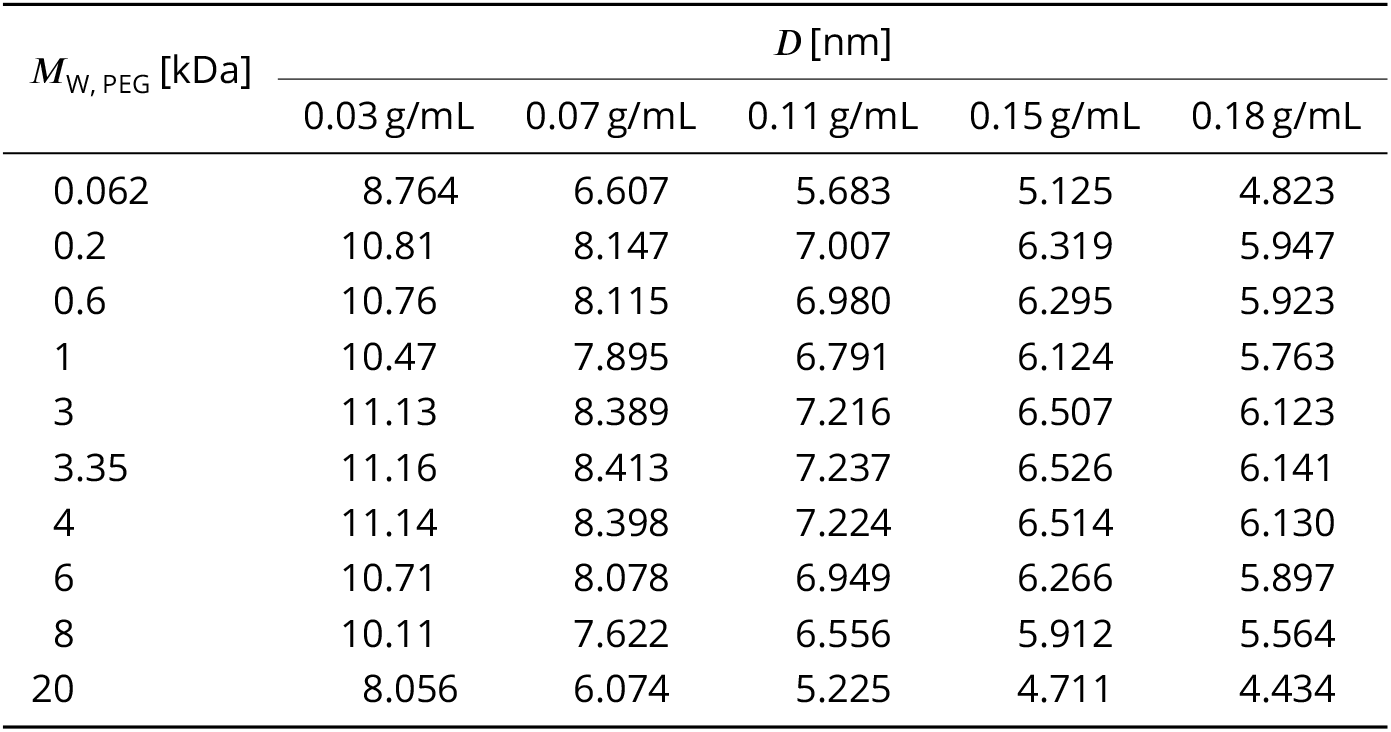
*D* of all used PEG variants at a concentration range of 0.03 to 0.18 g/mL.

**Figure 7—figure supplement 1.**
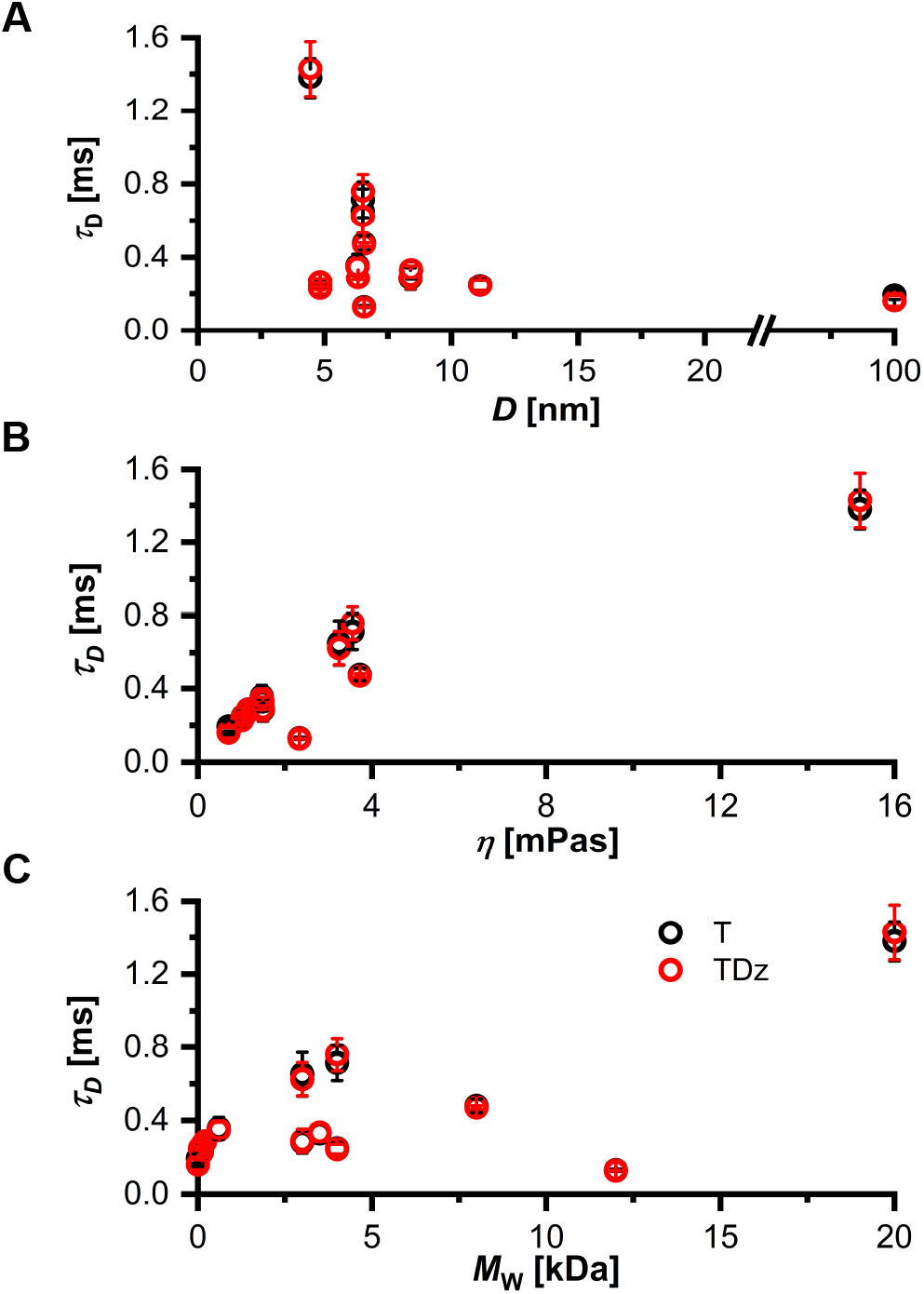
Diffusion time *τ*_D_ of RNA substrate (T) and Dz:RNA complex (TDz) in absence and presence of cosolutes determined by FCS.. **A** *τ*_D_ depending on cavity size *D*. **B** *τ*_D_ depending on the viscosity of crowded solutions determined at 37 °C. **C** *τ*_D_ depending on *M*_W_ of the respective cosolute.

**Figure 7—figure supplement 2.**
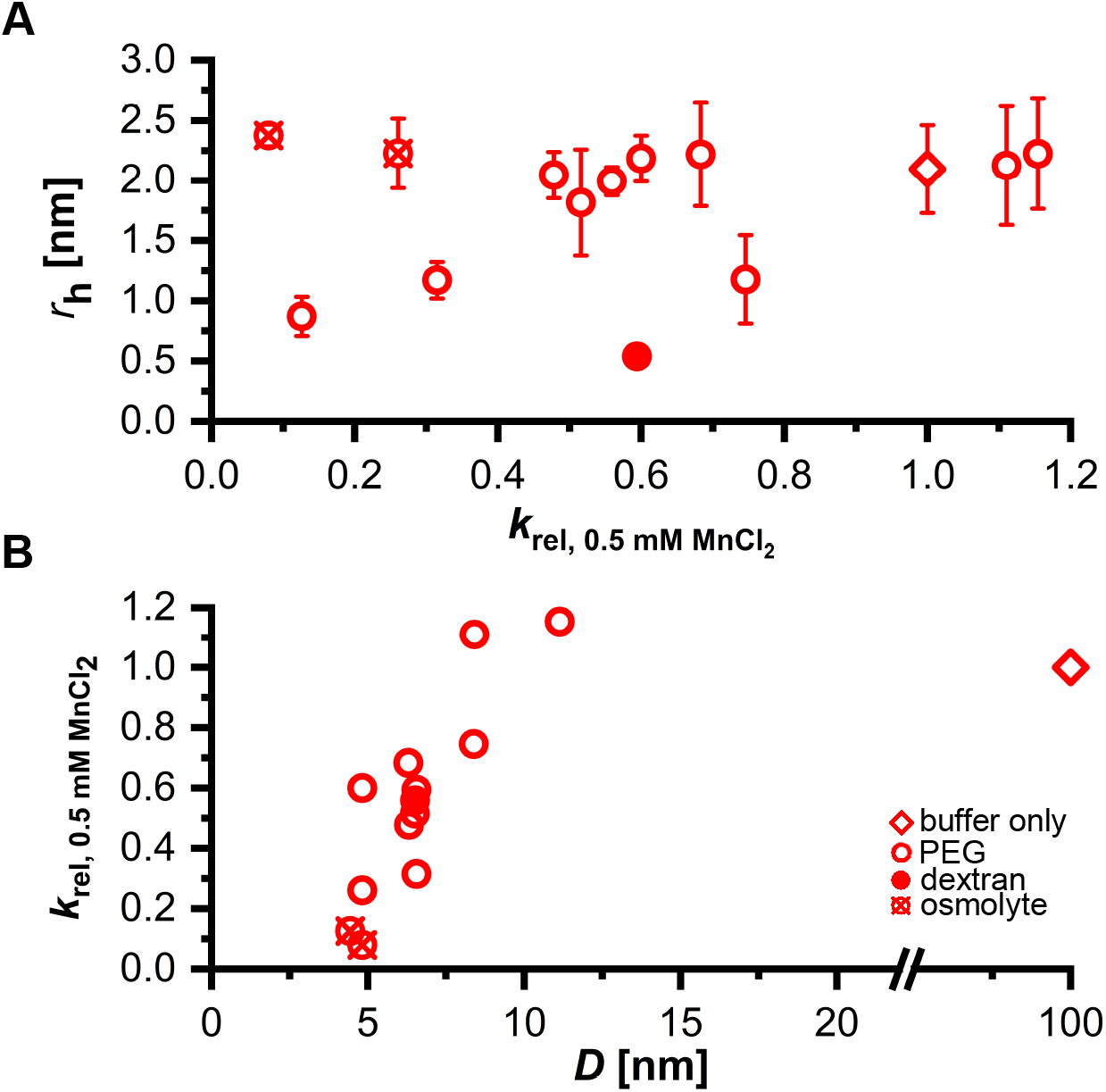
Comparison of the Dz:RNA complex’s hydrodynamic radius *r*_h_, *D*, and relative Dz activity for Mn^2+^-induced RNA cleavage.. **A** *r*_h_[nm] of TDz in comparison to *k*_rel_ at 0.5 mM MnCl_2_. **B** *k*_rel_ at 0.5 mM MnCl_2_ depending on cavity size *D*.

## Notes

### Competing Interest Statement

The authors have declared no competing interest.

